# Organelles harbour pH gradients

**DOI:** 10.64898/2025.12.12.694065

**Authors:** Sangyoon Lee, Sandip Chakraborty, Soyoung Kim, Asif Ali, Koushambi Mitra, Matthew Zajac, Anastasia Brown, David Pincus, Yamuna Krishnan

**Affiliations:** Department of Chemistry, The University of Chicago, Chicago, IL, USA; Neuroscience Institute, The University of Chicago, Chicago, IL, USA; Institute for Biophysical Dynamics, The University of Chicago, Chicago, IL, USA; Department of Molecular Genetics and Cell Biology, University of Chicago, Chicago, IL, USA; Center for Physics of Evolution, University of Chicago, Chicago, IL, USA

## Abstract

Organelle pH is critical to organelle identity and function. Resident proteins that define each organelle modify transiting cargo proteins, with both retention and trafficking between organelles governed by pH-dependent mechanisms. For example, lysosomal enzymes bind mannose-6-phosphate receptors at the higher pH (∼6.5) of the Golgi and dissociate at the lower pH (∼5.5) of late endosomes^1^. Proteins that stray from the endoplasmic reticulum (ER) are captured by KDEL receptors in the acidic Golgi and returned into the neutral ER^2,3^. This pH-tuned trafficking system compartmentalizes organelle function and prevents mis-localization of critical enzymes^4^. Dysregulated organelle pH disrupts their function and leads to various diseases. Because protons move rapidly in water, the pH within a single organelle is currently assumed to be spatially uniform^5^. Here, using a reporter sensitive from pH 5.5 - 10.5 to map a spectrum of organelles at high resolution, we discovered that pH gradients exist within single, large or long organelles such as the ER and mitochondria, and in membrane-less organelles without ion-transporting proteins such as the nucleolus. These new findings upend our understanding of organellar pH, prompting new questions about proton diffusion within the cell, and its potential consequences on organelle function.

---

Organelles are like chemical reactors arranged in series. Each organelle with its distinct set of resident proteins and lumenal pH modifies cargo proteins in a stepwise fashion as they move through the cell. To ensure the cargo is processed in time before the organelle matures, the catalytic rates of lumenal enzymes are aligned with organelle trafficking times. Therefore, enzyme activity is evolutionarily optimized to the organelle pH. Organelle pH in turn is sculpted by these lumenal metabolic reactions as well as active and passive ion transport across organelle membranes^4^. The transmembrane pH gradient in organelles provides a transducible electrochemical driving force. Organelles like lysosomes or mitochondria use this electrochemical driving force to move substrates unidirectionally^5^. Naturally, any dysregulation of organelle pH disrupts their function and leads to a panoply of diseases. For example, mutations in proteins that set mitochondrial pH, like UCP1, or those that use lumenal H^+^ to transport substrates, like SLC25A12/13, cause diabetes, fatty liver disease and encephalopathy^6–9^. Pathogens manipulate organelle pH to survive because phagosome acidification triggers their clearance^10^. Mutations in organellar Na^+^/H^+^ exchangers that regulate the pH of early endosomes, recycling endosomes, and the Golgi are respectively linked to autism, glioblastoma and mental retardation^11^. There is a potent connection between the loss of acidity in lysosomes and lysosomal storage disorders^12,13^. Restoring aberrant organelle pH could potentially treat diseases in humans^14^.

While we have identified the proton pumps, ion channels and transporters that help set and maintain organellar pH, a deeper and more nuanced understanding of how pH is regulated and shapes organellar function is still needed^5^. We currently assume that the pH within a single organelle is spatially uniform even though it is well known that different organelles of the same type can have different pH levels^14^ and that the pH of a single organelle can change over time^15^. This is because the small size of organelles and rapid diffusion of protons in water automatically implies that organelle pH must equilibrate instantly when protons enter or exit such closed systems. Yet there are two anomalous examples of sub-organellar pH gradients. Recently, linear and dynamic pH gradients were seen inside tubular lysosomes^16^, while a radial, yet stable, pH gradient was seen inside xenopus oocyte nucleoli^17^. Given that tubular lysosomes and nucleoli are very different structurally and functionally, and the pH gradients they harbour are also very different in terms of dynamics, length scales and acidity regimes, we posited that organellar pH gradients may be more prevalent than previously considered.

To test the above hypothesis, we need a way to map the pH of a wide range of organelles with a common molecular platform that is sensitive enough to capture sub-organellar pH differences if any. Because organelles span a wide range of pH, both anomalous examples of sub-organellar pH gradients were obtained with probes that were engineered specifically for those compartments and cannot be deployed widely. DNA-based probes that mapped pH in tubular lysosomes^14^ can be used only in endocytic organelles and cannot probe mitochondria or nucleoli. pH sensitive green fluorescence protein (GFP) variants such as pHluorin can be targeted to organelles, but its pH response and dynamic ranges render it incompatible with lysosomes. Further, pHluorins have shown inconsistencies with other probes^18^. Peroxisomes, which are supposed to be near-neutral, were reported to be mildly acidic when using a pHluorin probe^19^ suited for pH < 7.5, yet mildly basic when using a SNAFL-2 probe^20^ suited for pH > 7.2. However, pHluorin2 has enabled sub-organellar mapping of Golgi and nucleolar pH^21^. But while pHluorin2 revealed that the nucleolus is more acidic than the nucleoplasm, which is consistent with findings based on Hill-type probes and SNARF-4, these latter studies failed to note sub-nucleolar pH gradients^22,23^. Hence, a common pH mapping platform that targets an array of organelles, with an expanded coverage of pH and high sensitivity will both cross-verify the existence of sub-nucleolar pH gradients and reveal whether sub-organellar pH gradients are a broader phenomenon.

## A broad chemigenetic pH reporter

Our chemigenetic pH reporter, denoted SeRapHin, uses a fluorophore with an expanded pH-responsive regime and is based on a pentacyclic pyridinium scaffold ^24–26^. SeRapHin contains a ligand that targets organelles using a Halotag approach^27^ (**Supplementary Scheme 1 – 5**, **Supplementary Note 1 – 5**, and **Supplementary Fig. S1 – S5**). The expanded pH response arises from two ionizable, resonant phenolates (pKa_1_ = 7.4, pKa_2_ = 8.6) that generate three protonation states. Two of these – the doubly protonated (DP) and doubly deprotonated (DD) states – fluoresce strongly at distinct wavelengths (**Fig**. **1a**). DP has a donor-acceptor-donor system that fluoresces green (G) at acidic pH while DD has an extended donor-acceptor system that fluoresces red (R) at basic pH (**Fig. 1a**, **Supplementary Fig. S6**, and **Supplementary Note 6**). The change in the relative abundance of DP and DD states as a function of pH alters the ratio of G to R fluorescence (**Fig. 1b**). When DP dominates, G/R ratios sensitively report between pH 5.5 – 7.5 and when DD dominates, R/G ratios report between pH 7.5 – 10.5 (**Fig. 1c**). SeRapHin is bright enough to map pH in a wide range of organelles. When SeRapHin with a biotin moiety (SeRapHin^bio^) was immobilized on 100 nm streptavidin beads and imaged in G and R channels (**Methods**, **Supplementary Scheme 6**, **Supplementary Note 7**, and **Supplementary Fig. S7**), an approximately 20-fold change in either G/R or R/G values across the pH responsive range was observed (**Fig. 1d-f**).

**Figure 1.**
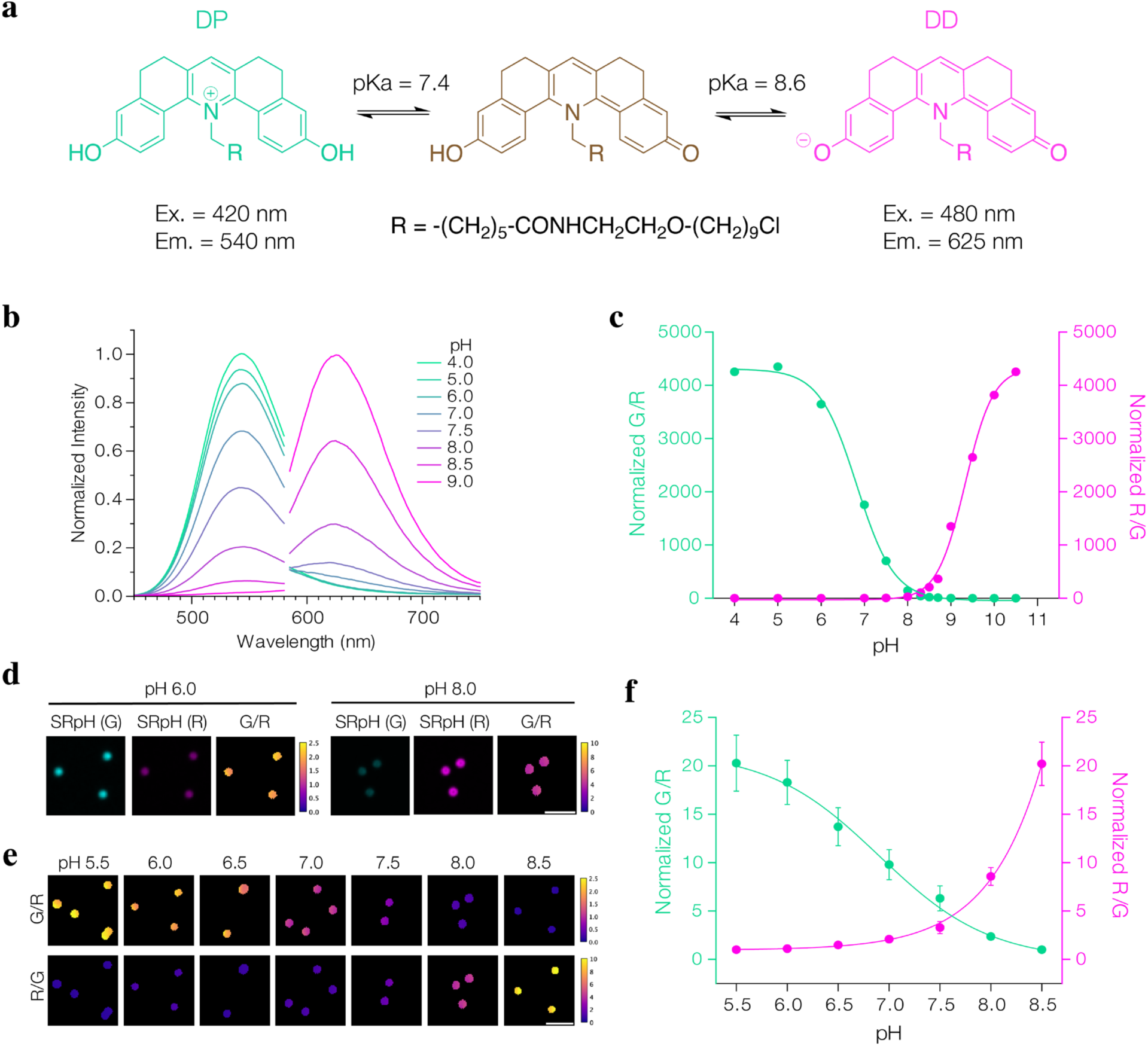
SeRapHin is a self-ratiometric pH reporter. **a**, Structure, pKa and fluorescence properties of the protonated states of SeRapHin. Doubly protonated (DP) and doubly deprotonated (DD) states emit green and red fluorescence, respectively. **b**, Fluorescence spectra of SeRapHin (G, λ_ex_= 420 nm and R, λ_ex_= 480 nm) shows decreasing G and increasing R intensities as pH increases. **c**, pH dependence of G/R (mint) and R/G (magenta) ratios of emission maxima (G, λ_em_= 540 nm and R, λ_em_= 625 nm) indicate an extended pH responsive regime. **d**, Confocal microscopy images show SeRapHin-coated beads display a clear pH response in the G (λ_ex_ = 445 nm, λ_em_ = 500 – 550 nm) and R (λ_ex_ = 488 nm, λ_em_ = 625 – 750 nm) channels at pH 6.0 and 8.0. **e**, Both G/R and R/G maps show SeRapHin-coated beads are responsive from pH 5.5 to pH 8.5 (bottom panels). Beads were 100 nm in diameter. Scale bar, 5μm. **f**, Bead-based pH calibration profile of SeRapHin over the full physiological regime. Plots are normalized either to G/R at pH 8.5 or R/G at pH 5.5. All experiments are in triplicate with n = 520 – 690 beads. Error bars are standard deviations of the means of each trial.

Because autofluorescence of cells can potentially interfere with or modify the probe readout, we calibrated the pH response of SeRapHin in cells by immobilizing it on the extracellular surface. We fused HaloTag7 (HT7) to the transmembrane domain of the PDGFRβ subunit bearing an IgK signal sequence^27^ and expressed this chimera, denoted HT7-PDGFRβ, in HeLa cells (**Fig. 2a**). The cells were then incubated with 1 µM SeRapHin for 2 h and thereafter washed twice with phosphate buffered saline (PBS) and once with pH buffer set at the target pH to remove unconjugated dye. Labeled cells were imaged in buffers ranging from pH 5.5 to 8.5 in G and R channels using a confocal microscope and ratio maps were constructed (**Fig**. **2b – d**, **Methods, Supplementary Note 8**, and **Extended Data Fig. 1**). Both G/R and R/G calibration curves as a function of pH in cells were consistent with SeRapHin response on streptavidin beads (**Fig. 2e – g**). The R/G curve fitted a sigmoidal curve across pH 5.5-8.5 (R^2^ = 0.9995), with a sharp increase at pH > 7. Similarly, the G/R curve followed a sigmoidal fit (R^2^ = 0.9975). The G/R fluorescence ratio is best for mapping mildly acidic to near-neutral organelles, whereas the R/G ratio is better for near-neutral to basic organelles.

**Figure 2.**
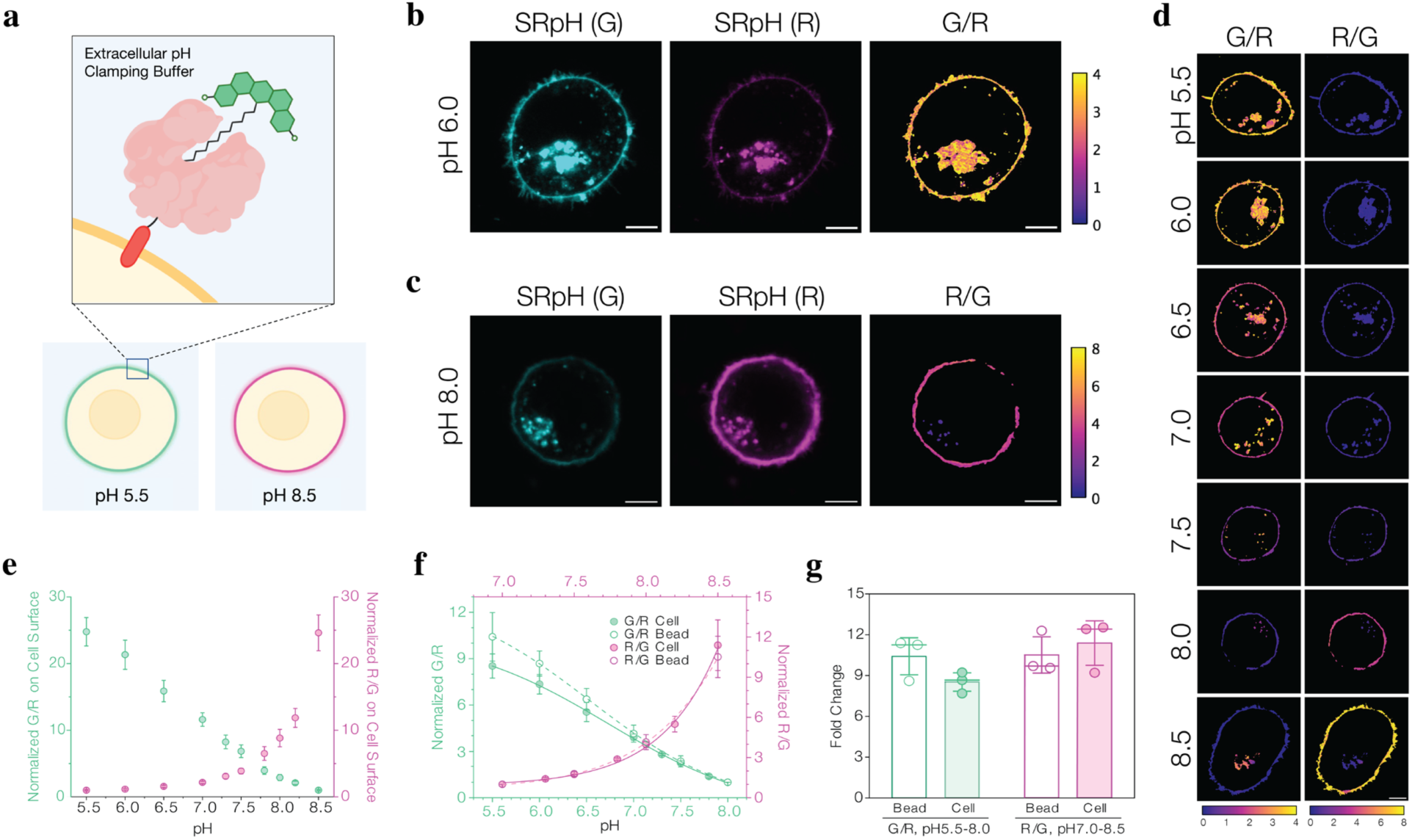
pH response of SeRapHin unchanged after conjugating to HaloTag domain expressed in cells. **a**, SeRapHin (green) is localized on the extracellular surface by reacting with HaloTag7 (HT7, salmon) expressed as a chimera with the transmembrane domain of PDGFRβ subunit (red). **b**, Confocal microscopy images showing SeRapHin’s pH response when immobilized on the cell surface in terms of G and R channel intensities at pH 6.0, and **c**, pH 8.0. **d**, G/R and R/G ratio maps as a function of pH for SeRapHin-labeled cells. **e**, Calibration curve of cell surface-immobilized SeRapHin as a function of pH obtained from heatmaps in **d**. G/R (mint) and R/G (magenta) are normalized to the values at pH 8.5 and pH 5.5, respectively. **f**, pH response of SeRapHin immobilized on cell surface (filled circles, solid lines) and beads (open circles, dashed lines) are similar in terms of G/R and R/G ratios. **g**, Fold-change of G/R (mint, from pH 5.5 to 8.0) and R/G (magenta, from pH 7.0 to 8.5) ratios for beads and cell surfaces in **f** are comparable. All experiments are in triplicate with n = 100 – 170 regions of interest. Error bars are standard deviations of the means of each trial. Scale bar in **b** – **d**, 5 µm.

### Mapping near-neutral to basic organelles

Given SeRapHin is sensitive from pH 6.0 – 8.5, we used it to probe neutral to basic organelles such as the ER, mitochondria and peroxisomes, which perform diverse chemistries such as peptide, ATP and plasmalogen synthesis. We targeted SeRapHin to these organelles in HeLa cells using previously described chimeras of HT7 and protein motifs (**Fig. 3a**)^27^. Specifically, to target the mitochondrial matrix, we fused HT7 to two repeats of the N-terminal Cox8a sequence (Mitomatrix-HT7)^27^. For peroxisomes, we fused HT7 to the PTS1 signal sequence (HT7-PTS1) and for the ER, we fused the calreticulin signal sequence to the N-terminus of HT7 and a KDEL motif to its C-terminus (ER-HT7). We tested the targeting specificity of these strategies by assessing the colocalization of the HT7 chimeras with key organelle markers. Mitochondrial targeting of Mitomatrix-HT7 was assessed by co-expression with the matrix marker, Mito-mScarlet^28^ (**Fig. 3b**). Peroxisome targeting was assayed by co-immunofluorescence of α-Pex14-Alexa647 and HaloTMR-labeled HT7-PTS1 (**Fig. 3c**). For ER targeting, we co-expressed Sec61β-GFP and ER-HT7 in cells and labeled the HT7 domain with HaloTMR (**Fig. 3d**).

**Figure 3.**
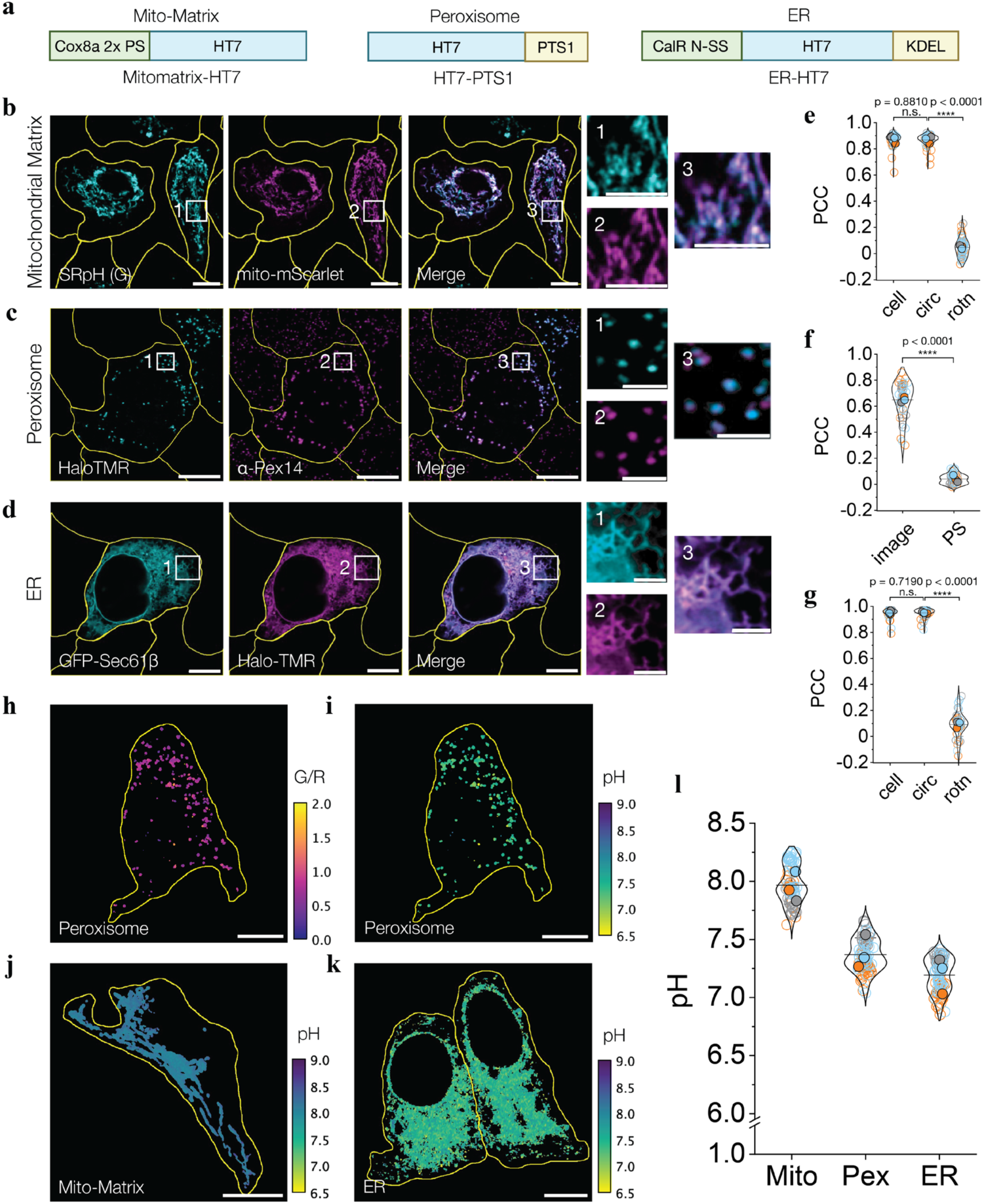
SeRapHin maps pH of near-neutral and basic organelles. **a**, HT7 chimeras for mitochondrial matrix (Mito-matrix), peroxisome, and ER. Cox8a 2x PS = 2 repeats of the Cox8a presequence; CalR N-SS = N-terminal signal sequence of calreticulin. **b-d**, Representative confocal images show SeRapHin-labeled Mitomatrix-HT7 colocalized with mitochondrial marker Mito-mScarlet (**b**), HaloTMR-labeled HT7-PTS1 colocalized with peroxisome marker α-Pex14(Alexafluor647) (**c**) and ER marker GFP-Sec61β colocalized with HaloTMR-labeled ER-HT7 (**d**). Merged images are on right and squares 1-3 are magnified on far right. **e-g**, Pearson’s correlation coefficient (PCC) of colocalization for **b**-**d** show robust organelle targeting (n_mito_ = 51, n_perox_ = 38, n_ER_ = 41). PS = pixel shift; rotn = image rotation (see **Supplementary Note 9**). Values are mean ± S.D. **h**, **i**, Representative G/R ratio map of peroxisomes in a single cell (**h**) and the corresponding pH map (**i**) derived following workflow in **Extended Data** Fig. 2. **j**, **k**, Representative pH map of mitochondria obtained from R/G map of one cell (**j**) and of ER obtained from G/R map of two adjacent cells (**k**). **l**, Mean pH values of mitochondrial matrix (Mito), peroxisome (Pex), and ER reported by SeRapHin. Each datapoint is the mean pH of the segmented organelle(s) in a single cell. All experiments are in triplicate. Each trial is colour coded. Mean value of each trial is in filled circle^57^ (n_mito_=76, n_perox_=88, n_ER_=91). n = number of cells. All scale bars, 10 μm, Magnified scale bars, 5 μm (mito-matrix), 2 μm (peroxisome), 3 μm (ER). Yellow lines in **b-d**, **h-k** outline cells.

Pearson’s correlation coefficient (PCC) for colocalization between SeRapHin (G channel) and Mito-mScarlet was 0.87 ± 0.05 (**Fig. 3e**, and **Supplementary Note 9**), that for peroxisomes was 0.65 ± 0.15 (**Fig. 3f**), and that of ER was 0.95 ± 0.05 (**Fig. 3g**), indicating robust targeting of these organelles. Following this, we used SeRapHin to map the pH of the ER, mitochondria, and peroxisomes (**Fig. 3h – k**, **Extended Data Fig. 2**, and **Supplementary Note 10, 11)**. Consistent with previous reports^18,29,30^, the mean mitochondrial matrix pH reported by SeRapHin was 7.97 ± 0.02 and that of the ER was pH 7.19 ± 0.02 (**Fig. 3l**, and **Extended data Fig. 3**). The average pH of peroxisomes, however, was 7.37 ± 0.02, a finding we will discuss in greater detail below (**Fig. 3l**).

## SeRapHin resolves sub-organellar pH

To test whether SeRapHin can resolve pH differences at sub-organellar resolution, we mapped the well-known pH gradient along the *cis*-Golgi, medial-Golgi, and *trans*-Golgi network (TGN) using HT7 chimeras Stx5-HT7, MGAT2-HT7, and B4GalT-HT7^27,31,32^ as targets, respectively (**Fig. 4a**). Syntaxin-5 (Stx5) is a component of the ER to Golgi t-SNARE complex in the cis-Golgi^33^, MGAT2-GFP localizes in the medial-Golgi with long retention times^34^ and B4GalT1^27^ is specific to the TGN^31,32^. Because it is difficult to localize proteins with sub-organellar resolution in the flat-packed architecture of native Golgi, we fragmented the Golgi into ministacks using nocodazole (**Methods**). This approach preserves the stacking order and enables the cis-, medial-, and trans-Golgi to be immunolabeled with spectrally distinct fluorophores. The resulting trilayer ministack reveals the precise localization of proteins within the Golgi subdomains^35^. Our markers for the cis-, medial- and trans-Golgi were antibodies against GM130, Giantin, or GalT-GFP, respectively.

**Figure 4.**
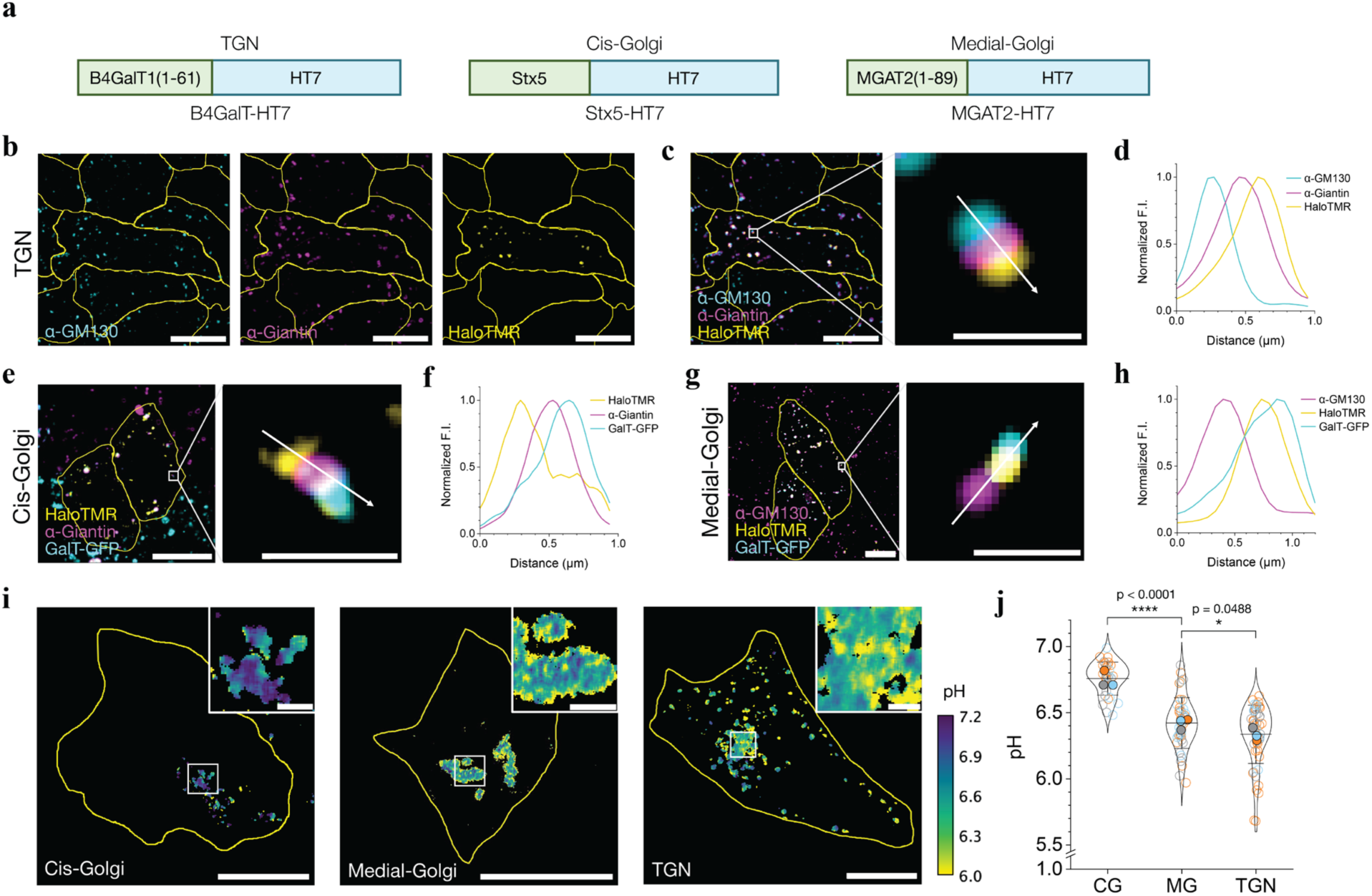
SeRapHin maps Golgi pH gradient at sub-organellar resolution. **a**, HT7 chimeras targeting the trans-Golgi network (TGN), cis-Golgi, and medial-Golgi. B4GalT1(1-61), Syntaxin5 (Stx5), and MGAT2(1-89) are fused to the N-terminus of HT7. **b-h**, Representative confocal images of nocodazole-treated cells show our HT7 chimera localizes in the Golgi regions with high specificity. Localization of B4GalT-HT7 in TGN as reported by HaloTMR, and cis- and medial-Golgi by anti-GM130 and anti-Giantin, respectively (**b**). Merged image showing HaloTMR labeled B4-GalT-HT7 in the TGN in relation to cis- and medial-Golgi markers. (**c**). Line profile of fluorescence intensity in **c** showing relative localization of markers (**d**). Cis-Golgi localization of HaloTMR labeled Stx5-HT7 in relation to medial-Golgi marker (α-Giantin) and TGN (GalT-GFP) (**e**). Line profile of fluorescence intensity in **e** showing relative localization (**f**). Medial-Golgi localization of HaloTMR labeled MGAT2 in relation to cis-Golgi marker (α-GM130) and TGN (GalT-GFP) (**g**) Line profile of fluorescence intensity in **g** showing relative localization (**h**). Merged and magnified images, and line intensity profiles show a well-resolved Golgi ministack in all cases. **i**, pH maps of cis-Golgi (left), medial-Golgi (middle), and TGN (right) obtained using SeRapHin. Insets: magnified views of the Golgi compartments. **j**, Mean pH values of the cis-Golgi (CG), medial-Golgi (MG) and TGN. Each datapoint is the pixel mean pH of the segmented organelle(s) in a whole cell. (n_CG_=31, n_MG_=46, n_TGN_=52). All experiments are in triplicate. Data from each trial is colour coded. Mean value of each trial is in filled circle of the corresponding colour^57^. All scale bars, 10 μm. All scale bars for magnified view, 1 μm.

We probed the specificity of localization of our HT7 chimera by labeling with HaloTMR. So, to test the TGN targeting specificity of B4GalT-HT7, we labeled it with HaloTMR and labeled the medial- and cis-Golgi with α-Giantin and α-GM130 markers, respectively (**Fig. 4b**). This yields a clear trilayer morphology, where Halo-TMR was adjacent to, but did not colocalize with α-Giantin, which in turn, was adjacent to, but did not colocalize with α-GM130. (**Fig. 4c, d**) Similarly, Stx5-HT7 and MGAT2-HT7 localized clearly in the cis- and medial-Golgi stacks, respectively (**Fig. 4e – h**, **Supplementary Fig. S8**, and **Supplementary Note 12**). Mapping the pH of these distinct regions revealed a pH of 6.75 ± 0.02 for cis-Golgi, 6.42 ± 0.03 for medial-Golgi, and 6.34 ± 0.03 for TGN (**Fig. 4i, j**). The TGN value is higher than those reported previously^14,36^ (pH 6.18 ± 0.01) because B4GalT-HT7 unavoidably leaks out of the TGN (**Supplementary Note 10**). For studying TGN, retrogradely trafficking probes^14,37,38^ are a better option. Nevertheless, SeRapHin clearly resolves the pH gradient in the Golgi at sub-organellar resolution.

When used to probe peroxisomes, SeRapHin revealed that peroxisomal pH spans a broader range than previously thought. Earlier measures yielded a pH of 6.9–7.1 while a recent measure yielded a pH of 7–8, suggesting that peroxisomes are more basic than neutral^19,39^. The high sensitivity of SeRapHin in both regimes shows that peroxisomal pH encompasses both extremes: pH 6.8 – 7.8 (**Fig**. **5a**-**b**, **Extended Data Fig. 4**). To test whether this variability in pH is genuine or a result of an imaging artifact, we fixed the cells in methanol and mapped the peroxisomal R/G values (**Supplementary Note 13**). Methanol treatment disrupts the organelle membrane and allows protons to equilibrate across compartments, effectively collapsing peroxisomal pH to cytosolic levels. Under these conditions, R/G values are expected to become uniform throughout the organelle. If imaging artifacts were responsible, R/G values would persist even after fixation. Our results show R/G ratios of fixed peroxisomes became uniform, indicating that the pH variability observed is genuine (**Fig. 5c**, and **Supplementary Fig. S9**).

**Figure 5.**
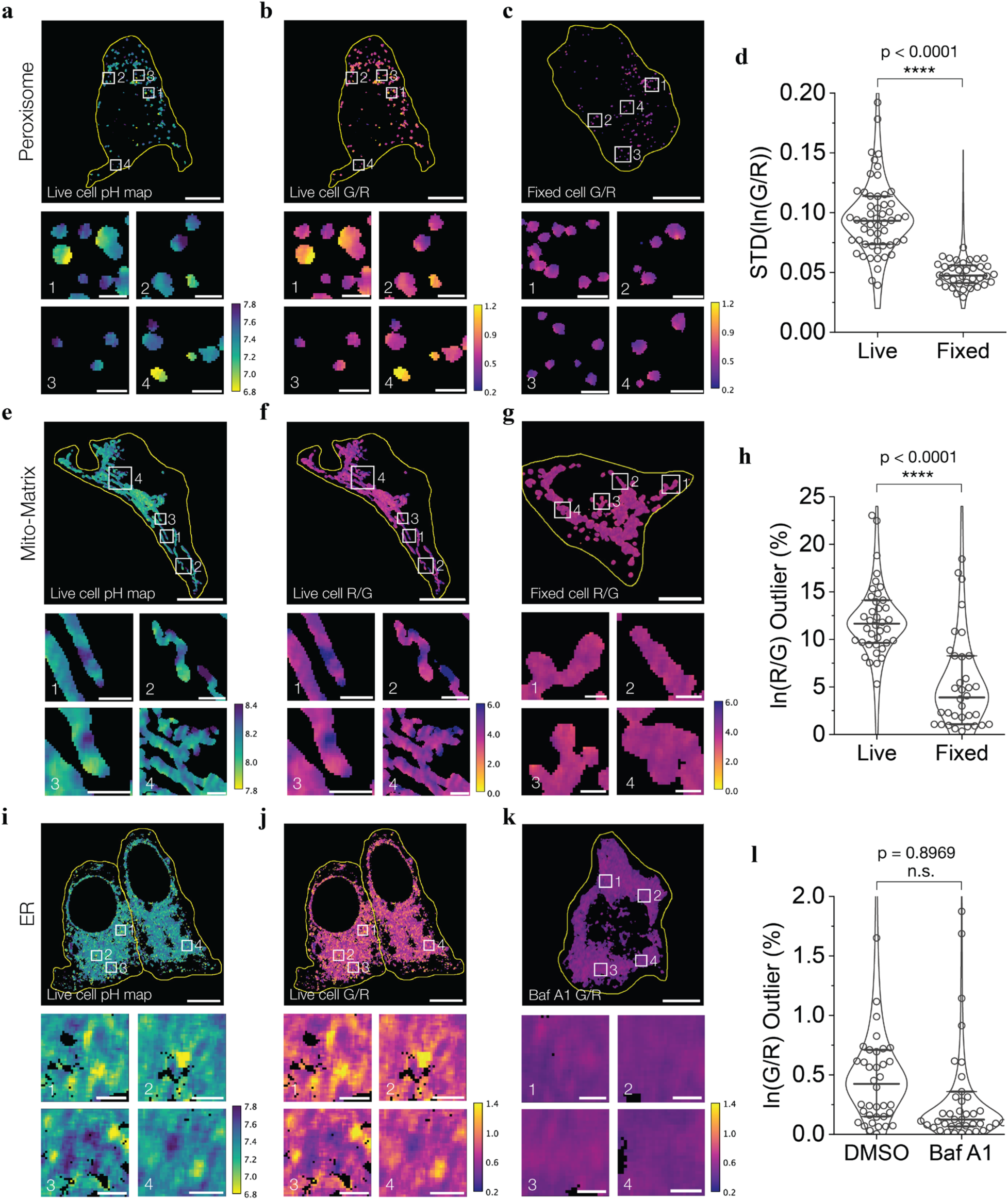
SeRapHin reveals pH gradients in individual peroxisomes, mitochondria and ER. **a**-**d**, Representative high-resolution pH (**a**) and raw G/R (**b, c**) heatmaps of SeRapHin-labeled peroxisomes in live HeLa cells (**a**, **b**) and those fixed with methanol (**c**). Heatmap (**c**) and quantitative measurements (**d**) show collapsing peroxisomal pH through methanol fixation led to monodisperse G/R values (**c**) and lower pH variance (**d**). Each datapoint in **d** shows standard deviation of ln(G/R) of peroxisomes in a single cell. (n_live_ = 54, n_fixed_ = 36) **e-h**, Representative high-resolution pH (**e**) and raw R/G (**f, g**) heatmaps of SeRapHin-labeled mitochondria in live (**e**, **f**) and methanol-fixed (**g**) cells. Single mitochondria in live cells show pH variability (**e**, **f**). Fixation led to uniform R/G values (**g**) and collapsed pH variance (**h**). Each datapoint is the outlier pixel number ratio with high ln(R/G) values. (n_live_ = 38, n_fixed_ = 34) **i**-**l**, Representative high-resolution pH (**i**) and raw G/R (**j, k**) heatmaps of SeRapHin-labeled ER in live cells (**i**, **j**) and those fixed with Bafilomycin A1 (**k**). ER of a single live cell shows pH differences (**i, j**). Collapsed G/R values in the heatmap (**k**) and calculated percentage of outlier pixels (**l**) after Bafilomycin A1 treatment indicate pH variability seen in **i** is genuine. DMSO is vehicle-treated cells. (n_DMSO_ = 38, n_Baf A1_ = 42, extreme outliers (ln(G/R) > 2.0 % are not shown but included in statistical analysis (t-test, IQR)). All scale bars, 10 μm. All magnified scale bars, 1 μm. Yellow lines outline cell. Lines and whiskers in **d**, **h**, **l**, represent medians and interquartile ranges. Boxes 1-4 in **a-c**, **e-g**, and **i-k** are magnified below each figure panel.

We further verified the pH variability by comparing the variance and standard deviation of ln(G/R) values per cell for peroxisomes in fixed and live conditions (**Supplementary Note 14**). Because fixation collapses transmembrane pH gradients, fixed cells showed much lower standard deviation, corroborating the pH variation observed in live cells (**Fig. 5d**). These results reconcile the discrepancy between previous measurements in the literature. In those cases, each probe likely captured only the subset of peroxisomes within its optimal pH response regime. By mapping a broader pH window, we reveal the full spectrum of peroxisomal pH values and resolve probe-specific biases. Given peroxisomal pores are large enough for folded proteins to translocate through^40^, the presence of a transmembrane pH gradient across peroxisomes is somewhat surprising. However, recent work suggests that these pores open and close intermittently^41^. When pores are closed, proton-generating metabolic activities such as fatty acid chain oxidation or ion transport likely drive the buildup of a proton gradient within the peroxisomes. Pore opening likely resets the gradient.

## pH domains in large single organelles

Within large and complex single organelles like mitochondria and ER, high-resolution pH maps obtained using SeRapHin revealed domains of different pH. Although mitochondria tend to clump in HeLa cells, many still show high aspect ratios and are individually addressable. Examining the pH maps of such mitochondria, we found an approximately 4-fold variation in the R/G ratio, spanning pH 7.8 – 8.4, harbored within a single organelle (**Fig. 5e, f**, **Supplementary Fig. S10**, and **Supplementary Note 15**). To test if this pH variability is genuine, we repeated the experiment in fixed cells (**Supplementary Note 13**) and quantified the heterogeneity by calculating the fraction of outlier pixels, or pixels with ln(R/G) values higher than the average (**Supplementary Note 16**). The much lower ratio of outlier pixels when fixed indicates that the pH variability is a feature of native mitochondria (**Fig. 5g, h**). Our observation supports recent studies showing individual mitochondria cristae functioning independently with distinct membrane potentials^42^, aligns with evidence of restricted solute diffusion in the mitochondrial matrix^43^ and pH differences in the intermembrane space^44^.

In the case of ER, we observed a spatially non-uniform pH. Even though the pH was 7.21 ± 0.02 on average, there were regions in the ER that were as acidic as pH ∼6.8 (**Fig. 5i, j**). To test whether these hotspots of acidity are genuine pH differences, we mapped the pH after collapsing the gradient. Given the documented presence of V-ATPase at ER exit sites (ERES), we collapsed the gradient by blocking the proton pumping action of V-ATPase using Bafilomycin A1. Compared to vehicle control cells treated with dimethyl sulfoxide (DMSO), cells treated with Bafilomycin A1 showed a much narrower spread of G/R as reflected by the significantly lower fraction of outlier pixels (**Fig. 5k, l**, and **Supplementary Notes 13, 17**). These observations reaffirm that the acidic hotspots are a feature of native ER. Their sensitivity to V-ATPase inhibition further suggests that these acidic hotspots correspond to ERES, where local micro-ERphagy occurs. Under ER stress, misfolded proteins and the V0 subunit of V-ATPase at the ERES recruit the V1 subunit from the cytosol to form a complete V-ATPase, triggering ER-phagy.

## pH gradients in membrane-less organelles

Besides membrane-bound organelles, SeRapHin can also be used to study membrane-less organelles at sub-organellar resolution. We studied the nucleolus because it is compartmentalized into three well-defined regions: GC (granular component), DFC (dense fibrillar component) and FC (fibrillar center) (**Fig. 6a**). Each region has a distinct molecular composition, and many proteins localize in specific sub-nucleolar zones^45^. During their biogenesis, ribosome intermediates move radially outwards from the center of the nucleolus, crossing these three distinct regions. We targeted SeRapHin to each of these regions in U2OS human osteosarcoma cells using HT7 fusions of such localized proteins (**Fig. 6b**). To target the GC, we stably expressed the *bona fide* GC marker NPM1 with HT7 (HT7-NPM1) in U2OS cells^17^. For DFC, we targeted the FBL protein by stably expressing HT7-FBL and for the FC region, we expressed HT7-RPA43. (**Extended Data Fig. 5**) mNeongreen-FBL has been used to study FBL mobility^46^ and mCherry-RPA43 is known to localize selectively in the FC^47^. Unlike previous measures where the error is the standard error of the mean (s.e.m.), for membrane-less organelles we report standard deviation because the s.e.m. is in the order of 10^-3^.

**Figure 6.**
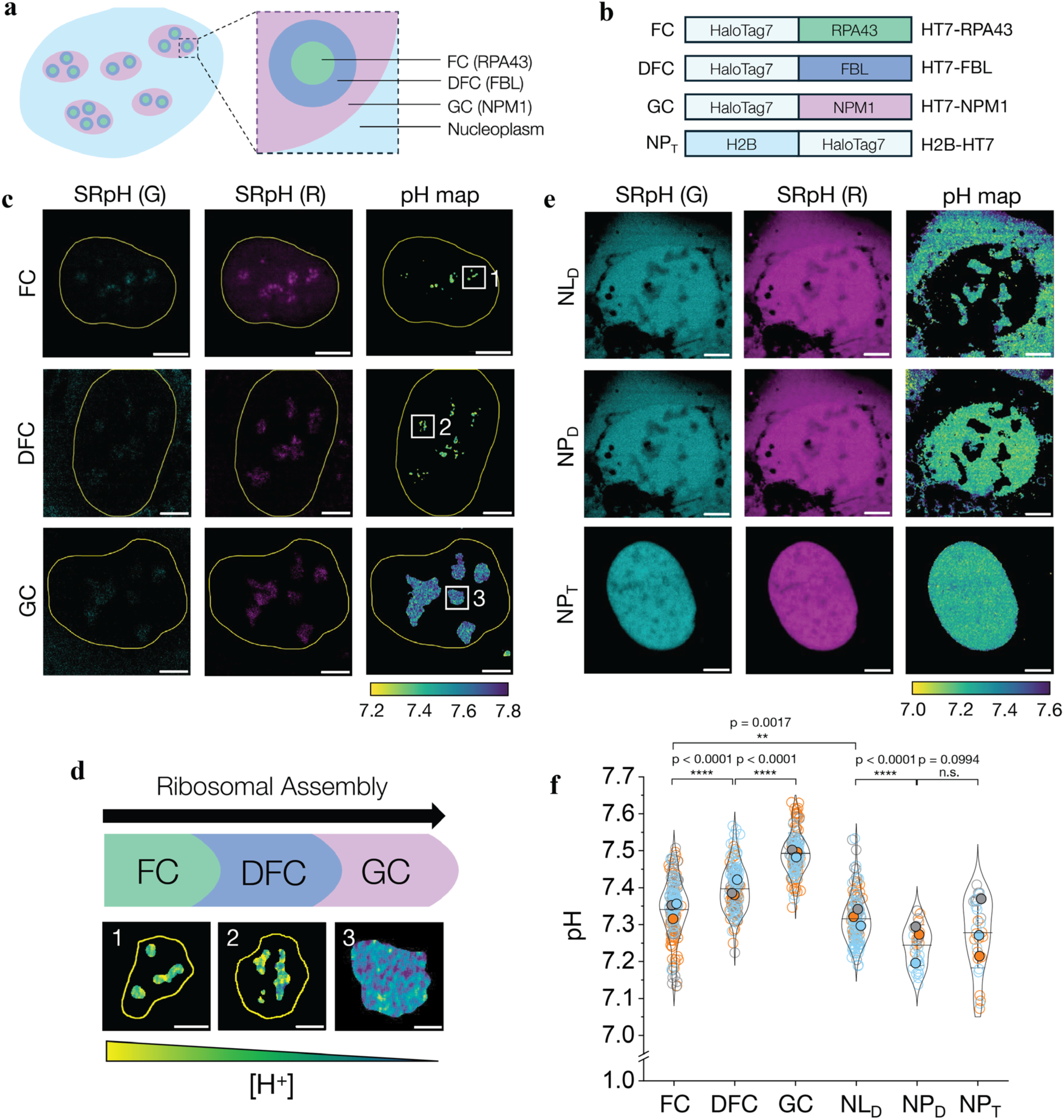
Membrane-less organelles like nucleolus harbour pH gradients. **a**, Nucleolus has three regions: fibrillar component (FC), dense fibrillar component (DFC), and granular component (GC). **b**, HT7 chimeras of RPA43, FBL, and NPM1 targeting SeRapHin to GC, FC and DFC regions of the nucleolus. H2B-HT7 targets SeRapHin to chromatin in the nucleoplasm (NP). **c**, Representative images in G (left column) and R (middle column) channels and the corresponding pH heatmaps (right column) of SeRapHin-labeled FC (top row), DFC (middle row) and GC (bottom row) regions of nucleoli within a single cell nucleus. White squares 1-3 are magnified in **d**. **d**, Reaction coordinate of ribosome biogenesis across the sub-nucleolar zones as a function of local pH. High-resolution magnified pH maps for each zone are from boxes 1-3 in **c**. **e**, Representative images in G (left column) and R (middle column) channels and the corresponding pH heatmaps (right column) of nucleolus (NL_D_) and nucleoplasm (NP_D_) reported by passively diffusing SeRapHin conjugated to HT7-PTS1, and of the nucleoplasm when SeRapHin is targeted to chromatin by H2B-HT7 (NP_T_). Image analysis described in **Extended Data** Fig. 2. **f**, Calculated mean pH values of nucleolus regions (FC, DFC, GC), nucleolus and nucleoplasm reported using a diffusing probe (NL_D_, NP_D_) and nucleoplasm reported using a targeted probe (NP_T_). All experiments are in triplicate. Data from each trial is colour coded. Mean value of each trial is in filled circle^57^ (n_FC_ = 165 n_DFC_ = 155 n_GC_ = 141 n_NLD_ = 146, n_NPD_ = 31, n_NP_ = 31, n = number of nucleoli / nuclei). Yellow lines outline nucleus (**c**) and nucleolus (**d**). Scale bars, 5 µm (**c**, **e**) and 1 µm (**d**).

Our measurements show a small but statistically significant pH gradient across FC (pH 7.34 ± 0.08), DFC (7.40 ± 0.06) and GC (7.49 ± 0.06) in the human cell nucleoli (**Fig. 6c, d, f**). A radial pH gradient but lacking sub-nucleolar specificity was previously observed in the nucleoli of xenopus oocytes and was posited to provide a proton motive force^17^. Interestingly, while previous qualitative maps of nucleolar pH in human cells revealed differences between the nucleoli and nucleoplasm, no sub-nucleolar pH gradients were evident^23^. This could be due to the probes used and/or architectural differences between human and xenopus nucleoli. Those prior studies used freely diffusible probes, which are not localized to a specific sub-nucleolar region. In oocytes, the nucleolus is a simple tri-layer structure composed of a DFC core jacketed by FC and then GC, whereas in humans, the nucleolus has multiple FC and DFC cores embedded in the GC (**Fig. 6a**).

To cross-verify the sub-nucleolar pH gradients seen in our experiments and understand why these features were missed earlier, we mapped the pH of the diffusible space within the nucleolus (NL_D_) and nucleus (NP_D_) of U2OS human cells using one type of HT7 chimera for both compartments. We chose HT7-PTS1 as the diffusible probe because while this chimera is translated in the cytoplasm and enriched in peroxisomes, it is also present in the nucleoplasm at low levels and in the nucleolus at even lower levels. Through intensity-based segmentation, we clearly resolved both the nucleus and nucleoli (**Fig. 6e**). As a control (NP_T_), we mapped the pH of the nucleus with a non-diffusible probe by incorporating SeRapHin into chromatin via HT7-H2B. Intriguingly, the space accessible to diffusing probes in the nucleolus spans the entire volume of the nucleolus and displays a more uniform pH (7.32 ± 0.06) that is distinct from, and slightly more acidic than the punctate FC, DFC and GC regions seen in **Fig**. **6c**. These observations suggest that when localized to a protein that is confined to a specific sub-nucleolar zone, SeRapHin reports the pH of that local, specialized microenvironment. The nuclear pH map obtained by targeting SeRapHin to chromatin, showed no spatial heterogeneity (**Fig. 6e**) and had an average pH (7.28 ± 0.10) that was consistent with that of the diffusible space, NP_D_ (7.24±0.06) (**Fig. 6f**) and other NP_D_ measurements reported previously^48,49^. These results collectively confirm that like xenopus oocytes, human nucleoli harbour sub-nucleolar pH gradients, and probe diffusion is likely one of the reasons why these features were missed previously.

It is interesting to consider how a membrane-less organelle like the nucleolus might harbour sub-organellar pH differences compared to the bulk. One explanation could be the dense, yet distinct, chemical composition of the sub-nucleolar zones. Nucleolar proteins in these zones interact with ribosomal RNA species that are at specific stages in their biogenesis. For example, NPM1 forms condensates with arginine-rich proteins like SURF6 and RPL23a that are also enriched in the GC^50^. Thus, NPM1 likely experiences a relatively basic environment due to the basic arginine-rich tracts of partner proteins in the condensate, and the high density of protonated arginines could provide a local electrostatic potential that would lower the local proton activity. This is chemically distinct from the environment of RPA43 in the FC that is depleted of most of the arginine-rich nucleolar proteins^17^. Another consideration is that protons are generated when nucleotides polymerize into ribosomal RNA. An actively transcribing, functional nucleolus produces ∼10^5^-10^6^ protons per second^51^. In an ideal, dilute aqueous solution, proton mobility is very high due to both diffusion of the protonated species and the Grotthuss mechanism^52–54^. The viscosity in the nucleolus, however, approaches ∼10 Pa.s, which is ∼10^4^ times that of water^55^. The co-existence of an active proton source and a sub-nucleolar pH gradient indicates that proton mobility is slower in this dense, crowded phase than in bulk solution.

## Discussion

The broad pH response of SeRapHin (pH 5.5-10.5) is due to two ionizable protons, unlike most pH indicators that have only one. The high dynamic range allowed us to capture local variations in pH across different organelles. When spatially averaged, the overall pH in each of our maps is consistent with previous organelle pH estimates. Besides probe diffusion, low dynamic range of probes and/or spatial averaging are other possible reasons for why sub-organellar pH gradients were overlooked previously. For example, despite a striking pH gradient in tubular lysosomes, their average pH is no different from vesicular ones^16^. Similarly in mitochondria and ER, the overall pH is consistent with previous measures^18^, but the high dynamic range of SeRapHin reveals ∼3-4 fold change in signal, corresponding to sub-organellar domains of lower and higher pH, thereby capturing previously muted spatial variations. In mitochondria, this heterogeneity, which is consistent with earlier reports that show individual cristae harbouring different membrane potential, implies distinct ionic milieus within the same organelle^42^. Prior reports of Ca²⁺ microdomains in the ER^56^ also suggest distinct ionic environments, which we have now extended to include pH, further supporting its architectural and functional heterogeneity.

For more precise delivery, next-generation SeRapHin probes would integrate its ultrasensitive, self-ratiometric readout with active targeting strategies. Such actively targeted probes would avoid collateral signals arising from basal autophagic flux or “biogenesis” intermediates produced in the ER. Alternatively, one can incorporate built-in strategies to mask unavoidable mistargeting due to autophagy or biogenesis. It is also critical to minimize structural perturbations associated with accommodating reporter protein loads in delicate locations such as the mitochondrial intermembrane space. With these advances, one can access the full spectrum of organelles, and enable the comprehensive mapping of sub-organellar pH at high-resolution.

We tend to consider aqueous pH as spatially constant, rarely considering protons as a moving species in water. Their speed by both diffusion and Grotthuss hopping has implied that the injection or ejection of protons in a closed system simply equilibrates pH instantly to a new value. Yet our data shows that inside the dense, crowded milieu of a cell, protons do not diffuse as they do in dilute aqueous solution. Their movement is sufficiently slow that local proton production or removal can create differential pH domains tens to hundreds of nanometres in size, prompting a reconsideration of the physicochemical constraints under which biological reactions operate. Precise measurements of viscosity in tubular or sheet-like ER, or mitochondrial matrix will complement our pH maps and enable better estimates of proton mobility in these environments. Such measurements are likely to be challenging because organelles are highly dynamic. Real-time maps of sub-organellar pH will reveal the lifetime and dynamics of these gradients and whether they guide or respond to architectural changes of the organelle. In the long term, uncovering how sub-organellar pH gradients shape organelle function may reveal new insights into disease etiology and offer new opportunities to harness pH modulation as a therapeutic strategy.

## Methods

### Reagents

^1^H NMR and ^13^C NMR spectra of the newly synthesized compounds were recorded on a Bruker AVANCE II+, 500 MHz NMR spectrophotometer in CDCl_3_ and tetramethylsilane (TMS) was used as an internal standard. Mass spectra were recorded with an Agilent 6224 Accurate-Mass time-of-flight (TOF) liquid chromatography–mass spectrometry (LC/MS). Streptavidin-coated microspheres were purchased from Bangs Laboratories, Inc., valinomycin and monensin were purchased from Cayman Chemicals. The antibodies used for immunofluorescence experiments in this study are as follows: α-PEX14(proteintech, 10594-1-AP), α-GM130(BD Biosciences, 610822), α-Giantin(abcam, ab80864), α-Giantin(abcam, ab37266), α-58K(abcam, ab27043), α-RPA194(F-6) (Santa Cruz Biotechnology, sc-46699), α-NOLC1(proteintech, 11815-1-AP), α-PES1(proteintech, 13553-1-AP), Goat anti-Mouse IgG (H+L), Alexa Fluor^TM^ 488(Invitrogen, A11001), Goat anti-Rabbit IgG (H+L), Alexa Fluor^TM^ 647(Invitrogen, A21245). HaloTag® TMR Ligand was purchased from Promega. All other reagents were purchased from Sigma-Aldrich (USA) unless otherwise specified.

### pH sensitivity of SeRapHin by fluorescence spectroscopy

100 nM of SeRapHin was aliquoted in UB buffer (10 mM HEPES, 10 mM MES, 140mM K^+^, 25 mM Na^+^, 1 mM Mg^2+^, 1 mM Ca^2+^, 20 mM OAc^-^, pH adjusted by HCl or KOH) Fluorescent emission spectra was taken by Fluoromax (Horiba) with the following collection parameters: For G fluorescence, λ_*ex*_ = 420 nm, range from 450 nm to 580 nm. For R fluorescence, λ_*ex*_ = 480 nm, range from 580 nm to 750 nm.

### SeRapHin^bio^ conjugation to streptavidin coated bead

1 µ m streptavidin coated polystyrene bead (Bangs Laboratories, CP01004) stock was first homogenized by vortexing. The 2 μL of bead stock was then dispersed into 15 μL of 20 μM SeRapHin^bio^ solution in deionized water. The mixture was then vortexed for 2 – 4 h while protected from light. After the incubation, it was centrifuged at 4 °C, > 16,000 g for 30 mins. Supernatant was discarded, and the collected beads are resuspended in 100 μL PBS. This washing step was repeated in total of three times. For the last resuspension, the beads were resuspended in 20 μL PBS to yield SeRapHin-coated bead stock solution, which was stored at 4 °C.

### SeRapHin imaging with confocal microscope

Leica Stellaris 8 was used for the pH imaging of SeRapHin. 63x objective (Leica HC PL APO CS2 63x 1.40NA) and Hybrid detectors (Leica HyD S, analog mode) were used. Line sequential scan is used, and pinhole was set to 95.5 μm (1AU for 580 nm). For SeRapHin (G), 448 nm laser with 15% intensity was used for excitation, and emission was collected from 500 nm to 550 nm (HyD S1 Gain 50 for beads, Gain 100 for cells). For SeRapHin (R), 488 nm laser was used for excitation (40% intensity) and emission was collected from 625 nm to 750 nm (HyD S3 Gain 50 for beads, Gain 100 for cells). In cases where DQ-BSA or Alexa Fluor 647^TM^ was used for lysosome masking, a third channel with 600 nm laser (5% intensity) and emission collected from 625 nm to 750 nm (HyD S3 Gain 20) was added.

### Bead calibration of the pH sensitivity of SeRapHin fluorescence

SeRapHin-coated bead stock solution was diluted in universal buffer (10 mM HEPES, 10 mM MES, 140 mM K^+^, 25 mM Na^+^, 1 mM Mg^2+^, 1 mM Ca^2+^, 20 mM OAc^-^, pH adjusted by HCl or KOH) with varying pH (from pH 5.5 to pH 8.5). The bead solution was drop casted on glass bottom dishes (Cellvis D35-10) and incubated in room temperature for > 20 mins to allow settlement to the bottom surface. Beads were imaged by confocal microscope (Leica Stellaris 8)

The single-plane bead images were analyzed in imageJ. Each channel was background subtracted, and bead pixels with significant signal intensity were segmented by Weka segmentation. Among the segmented beads, the ones that are solitary are selected as individual ROIs, and the mean intensity values from G and R channels were obtained to calculate the G/R or R/G of the bead. In the total triplicate data, more than 500 beads were analyzed per pH point. G/R values were normalized to the mean value from pH 8.5 as 1 for each replicate, and R/G values were normalized to the mean value from pH 5.5 as 1 for each replicate.

### Cell surface calibration of SeRapHin

HeLa cells of ∼70 % confluency seeded on glass bottom dishes (Cellvis 35-10) before more than 24 h were transfected with 250 ng of HT7-PDGFR, and incubated at 37°C, 5% CO_2_ for 18 – 24 h. Cells were then washed with PBS three times and treated with 1 μM SeRapHin in DMEM without FBS for 2 h at 37 °C, 5 % CO_2_. Cells are then washed with PBS three times and incubated in DMEM with 10% FBS at 37°C, 5% CO_2_ for over 4h. Cells are again washed with PBS two times and washed once with the pH buffer set to target pH. (10 mM HEPES, 10 mM MES, 140 mM K^+^, 25 mM Na^+^, 1 mM Mg^2+^, 1 mM Ca^2+^, 20 mM OAc^-^, pH adjusted by HCl or KOH). After adding the pH buffer again, the cells are imaged by confocal microscope (Leica Stellaris 8). The single plane G and R channel images are background subtracted and segmented via 2D Weka segmentation^58^. G/R and R/G ratio maps are constructed from the resulting images. ROI selection for obtaining the calibration curve is described in **Supplementary Note 8**.

### Colocalization experiments

Colocalization analysis is used to confirm SeRapHin targeting to specific organelle. For mitochondrial matrix, HeLa cells co-transfected with mitomatrix-HT7 and mito-mScarlet 18 h prior to the experiment was treated with SeRapHin following the protocol described in **Supplementary Note 10**. For peroxisome, HeLa cells transfected with HT7-PTS1 18 h prior to the experiment was treated with 5 μM HaloTag® TMR ligand in DMEM without FBS for 30 mins, washed with PBS three times and incubated in DMEM + 10 % FBS at 37 °C, 5 % CO_2_ for 1 h. Immunofluorescence for PEX14 was done following the method above. For ER, HeLa cells co-transfected with ER-HT7 and Sec61-GFP 18 h prior to the experiment was treated with 5 μM HaloTag® TMR ligand as described above. For TGN, HeLa cells transfected with B4GalT-HT7 18 h prior to the experiment was treated with 5 μM HaloTag® TMR ligand as described above. Then, the cells were treated with 33 μM nocodazole for 3 h. The cells are then immunolabeled with α-GM130 and α-Giantin. For cis-Golgi, HeLa cells co-transfected with Stx5-HT7 and GalT-GFP 18 h prior to the experiment was treated with 5 μM HaloTag® TMR ligand as described above. Then, the cells were treated with 33 μM nocodazole for 3 h. The cells are then immunolabeled with α-Giantin. For medial-Golgi, HeLa cells co-transfected with MGAT2-HT7 and GalT-GFP 18 h prior to the experiment was treated with 5 μM HaloTag® TMR ligand as described above. Then, the cells were treated with 33 μM nocodazole for 3 h. The cells are then immunolabeled with α-GM130. All samples were imaged under confocal microscope (Stellaris 8), and ImageJ plugin Coloc-2 was used for PCC analysis. The rotation method for testing the random colocalization and evaluating the colocalization is described in **Supplementary Note 9**.

### Immunofluorescence

HeLa cells on glass-bottom dishes (Cellvis 35-10) were washed three times with PBS and permeabilized with 0.2 % TritonX-100 in PBS and washed three times with PBS. Cells are then incubated in blocking buffer (3 % BSA in PBS) for 1 h. Blocking buffer was removed, and cells were incubated in primary antibody solution with 1% BSA for 1.5 h. The antibody dilutions are as follows: α-PEX14 (1:1000), α-RPA194 (1:200), α-NOLC1 (1:500), α-PES1 (1:100), α-GM130 (1:100), α-Giantin (rabbit, 1:1000), α-Giantin (mouse, 1:200), α-58K (1:200)). (0.25 % TritonX-100 + 0.1% Tween-20 in PBS for Golgi compartments) The cells were washed with PBS three times and incubated in secondary antibody solution with 1% BSA for 1 h. The antibody dilutions are as follows: goat-anti-rabbit IgG-AF647 (1:1000) and goat-anti-mouse IgG-AF488 (1:500). For Golgi compartments, PBST (0.1 % Tween-20 in PBS) was used instead of PBS for all washing steps after cell fixation. Blocking buffer and antibody solutions for Golgi compartments also included 0.1 % Tween-20. Then, the cells were washed with PBS three times and images at confocal microscope (Stellaris 8).

### Plasmids and cloning

HT7-PDGFRβ, MitoMatrix-HT7, HT7-PTS1, ER-HT7, B4GalT-HT7, and HT7-H2B were gift from Kai Johnson^27^, Mito-mScarlet was a gift from Gia Voeltz^28^. Sec61β-GFP was cloned from mCherry-Sec61β (gift from Christine Mayr)^59^, and CD-MPR-GFP-RUSH (gift from Juan Bonifacino) via Gibson assembly^60^. Stx5-HT7 was cloned from Stx5S-mCherry (gift from Geert van den Bogaart)^33^, and B4GalT-HT7 via Gibson assembly. MGAT2-HT7 was cloned from MGAT2-RpHLuorin2 (gift from Geert van den Bogaart)^21^, and B4GalT-HT7 via Gibson assembly.

### Cell culture

HeLa cells (ATCC, CCL-2) were grown according to the manufacturer’s protocol. In brief, Cells were grown in DMEM medium (Gibco, 11995) with 10% heat inactivated FBS (Gibco, 26140) and 100 U/mL Pen Strep (Gibco, 15140). Cells were maintained at 37 °C in a humidified chamber at 5% CO_2_ concentration. Cells are passaged with Trypsin-EDTA (0.25%) (Gibco, 25200072) five times diluted in PBS to be 0.05% trypsin. All the experiments were performed on cells at least 24 h after passaging. U2OS cells (ATCC, HTB-96) were grown with the same media and conditions as described with HeLa cells above. Cells are passaged with undiluted Trypsin-EDTA (0.25%) (Gibco, 2500072).

## Data analysis

Numerical data was processed and plotted with Origin software. For **Fig. 1f** and **Fig. 2e, f**, datapoint of each pH represents the average of G/R or R/G mean for each trial at the corresponding pH point, normalized to each trial’s pH point that gives smallest value of G/R or R/G. The standard deviation of the three trials is depicted by the error bar. For **Fig. 2g**, the fold-change of the G/R or R/G mean is depicted as a datapoint of an open circle. A median line is drawn with an error bar representing 1 S.D. to compare fold-change on beads and on cell surface. All colocalization and pH measurements in **Fig. 3-6** are represented as Superplots, where a PCC value from an ROI or mean pH value of the segmented organelle pixels from a cell is represented as an open circle data point that are color coded corresponding to the trial number. Different colors correspond to different trials. A filled circle was used to represent the mean of all PCC values or pH values for a particular trial^57^. Line represents the mean overall value of the combined data of the three trials. The error bars represent of 1 standard deviation. Unpaired t-tests with Welch’s correction were done to obtain two-tailed p values in **Fig. 4-6**.

## Data availability

The data supporting the figures in this paper will be uploaded onto a publicly available database post-acceptance and accession details will be specified here. All biological reagents will be available from the corresponding author upon reasonable request.

## Code availability

Code will be made available on a publicly available database post-acceptance, and accession details will be specified here.

## Acknowledgements

We thank Thomas R Cech, Ardem Patapoutian, Jack W. Szostak and Anthony A. Hyman for valuable discussions and critically reading the manuscript, as well as A.L. Chun of Science Storylab for critical input. Y.K. acknowledges NIH grants DP1GM149751, 1R01NS112139-01A1, 1R01GM147197-01, HFSP grant no: RGP0032/2022, the Bimla Rani Parkinson’s Disease Research Fund, the Ono Pharma Foundation. D. P acknowledges NIH grants R01 GM138689 and RM1 GM153533 and National Science Foundation QLCI QuBBE grant OMA-2121044.

## Author contributions

Y.K., S.L., S.C. and M.Z. conceptualized the study. S.L. and Y.K. designed the experiments. S.C. designed, synthesized and characterized SeRapHin and SeRapHin^bio^ *in vitro*. S.L. calibrated SeRapHin on beads and in cells. S.K. cloned MGAT2-HT7 and localized it in medial-Golgi. S.L. targeted and pH mapped all other organelles, with contributions from A.B. for the ER. S.L. optimized transfections, treatment times, and imaging with help from K.M. Y.K., D.P. A.A, and S.L. conceptualized and D.P. and A.A contributed reagents to, sub-nucleolar pH mapping. S.L. and A.A. selected and optimized imaging for HT7 fusions for nucleolar imaging. S.L., S.C. and Y.K. co-wrote the paper. All authors provided input on the manuscript.

## Competing interests

The authors declare no competing interests. YK is a co-founder of Esya Inc and MacroLogic Inc that use DNA nanodevices to develop diagnostics and therapeutics respectively.

## Additional information

### Supplementary information

SI Figures 1-4 and SI notes 1-9 are included in Supplementary Information file.

## Extended Data Figures

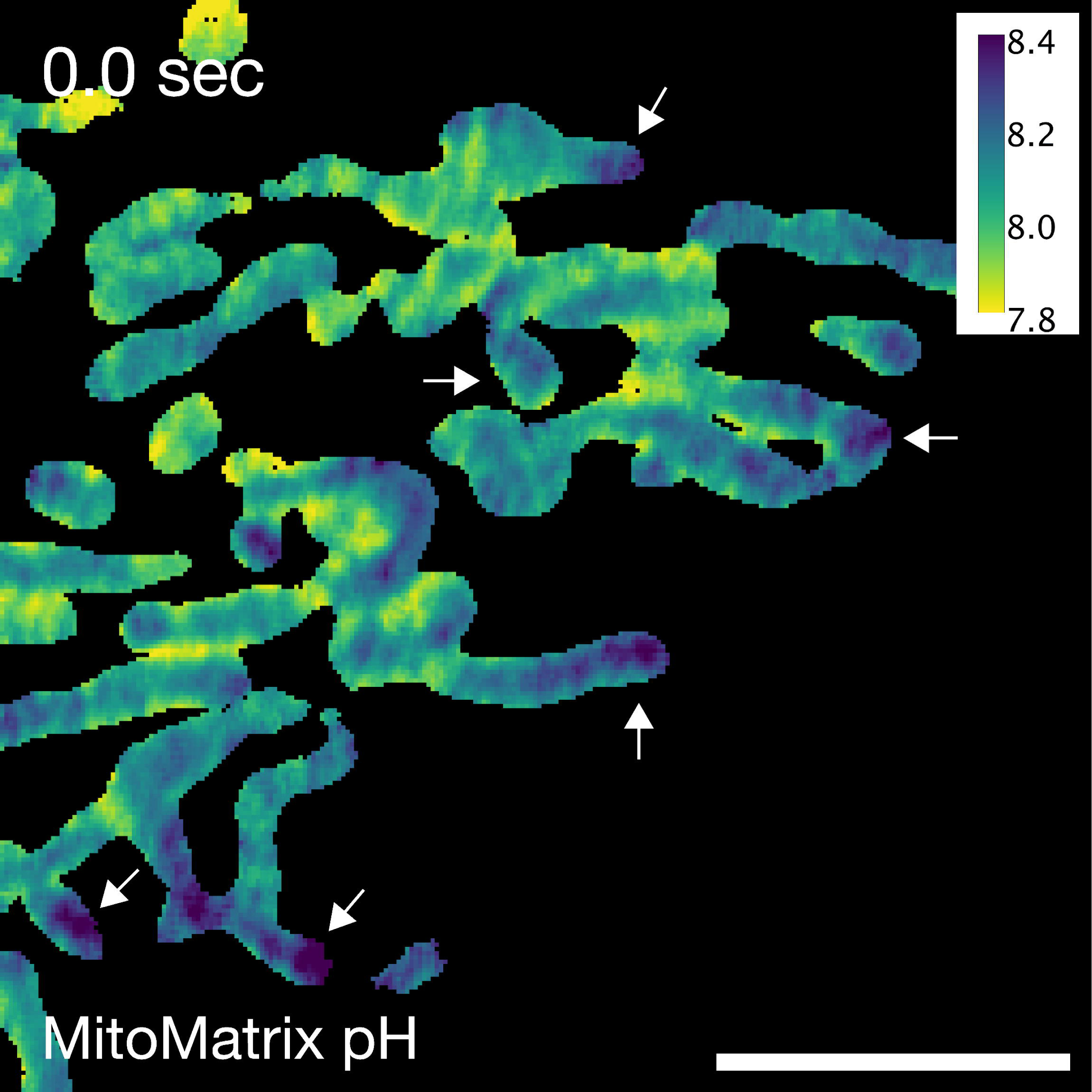

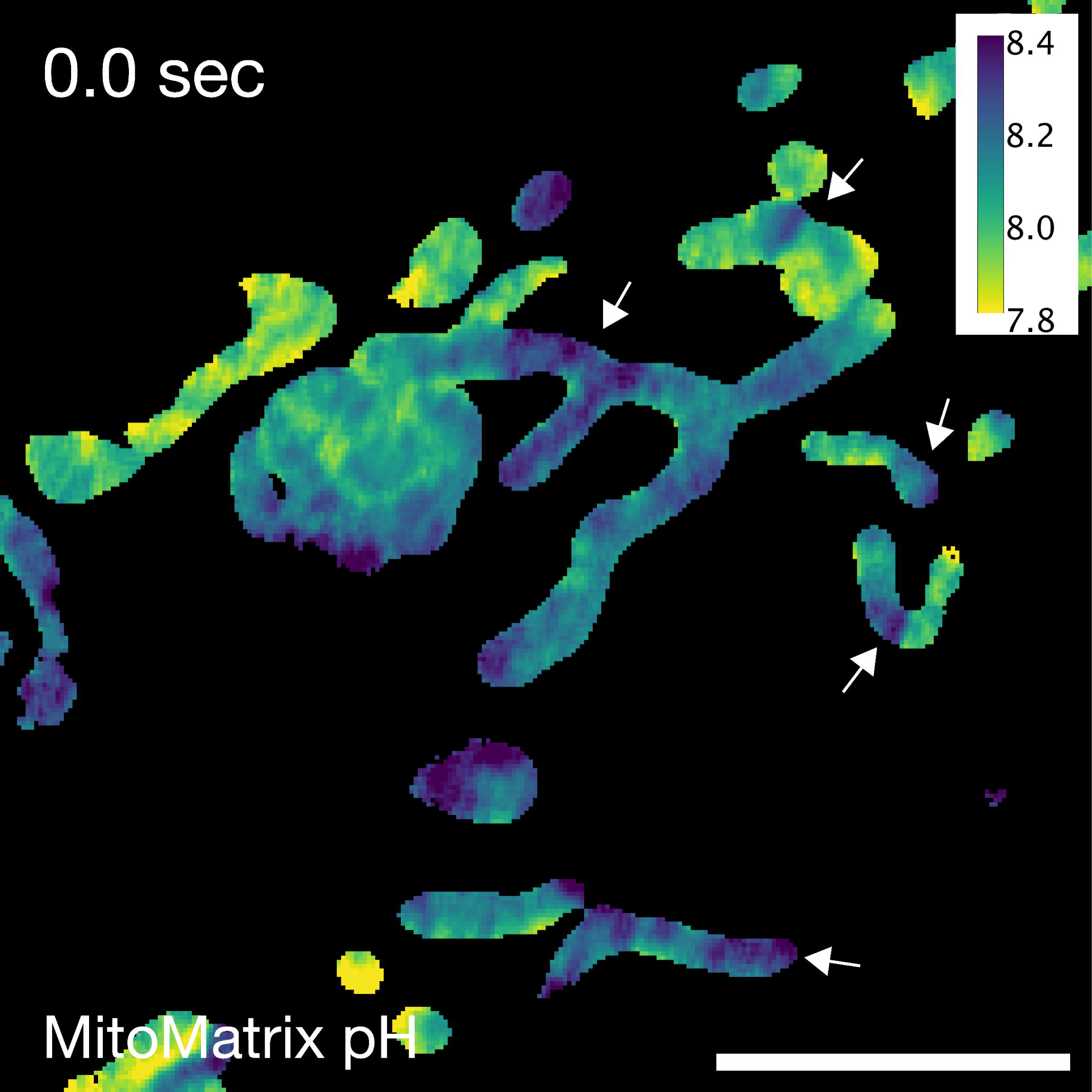

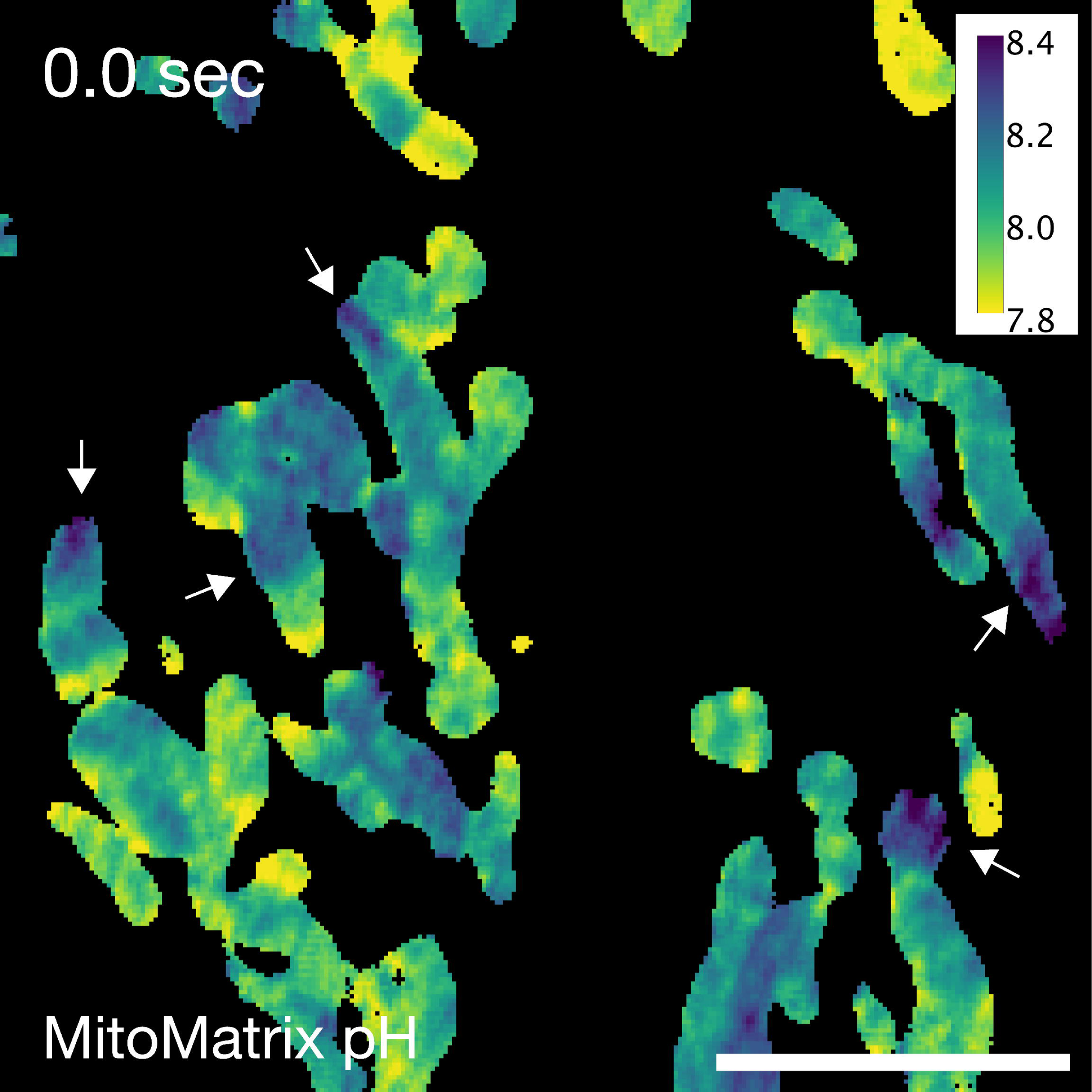

