## Extended Data Figures for "Organelles harbour pH gradients"

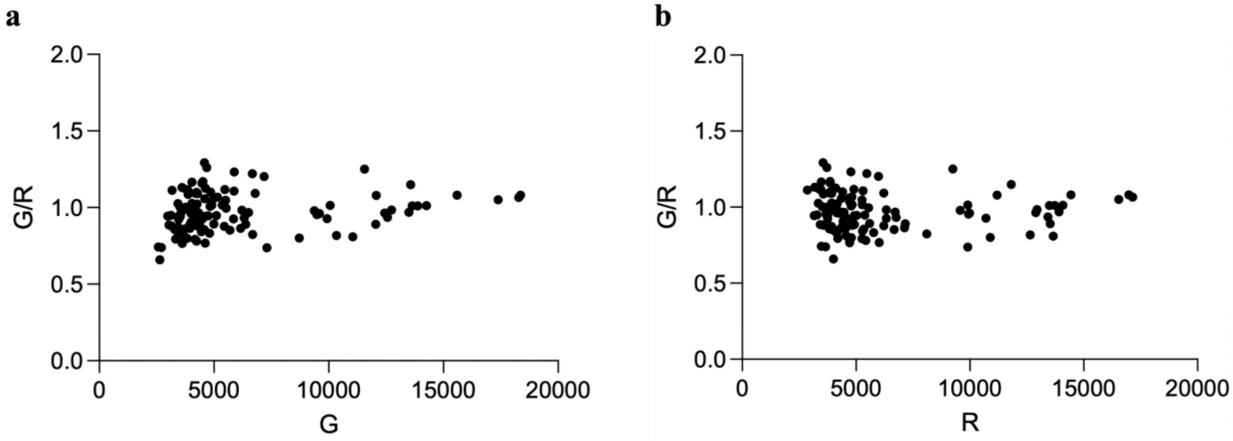


**Extended Data Fig. 1 | G/R and R/G values are independent of the overall SeRapHin signal intensity. a**, G/R – G plot of individual ROIs from the cell surface calibration image at pH 7.3 (**Methods**) shows that there is no significant dependence between G/R and G value. **b**, G/R – R plot of individual ROIs from the cell surface calibration image at pH 7.3 shows that there is no significant dependence between G/R and R value. (n = 121 ROIs)

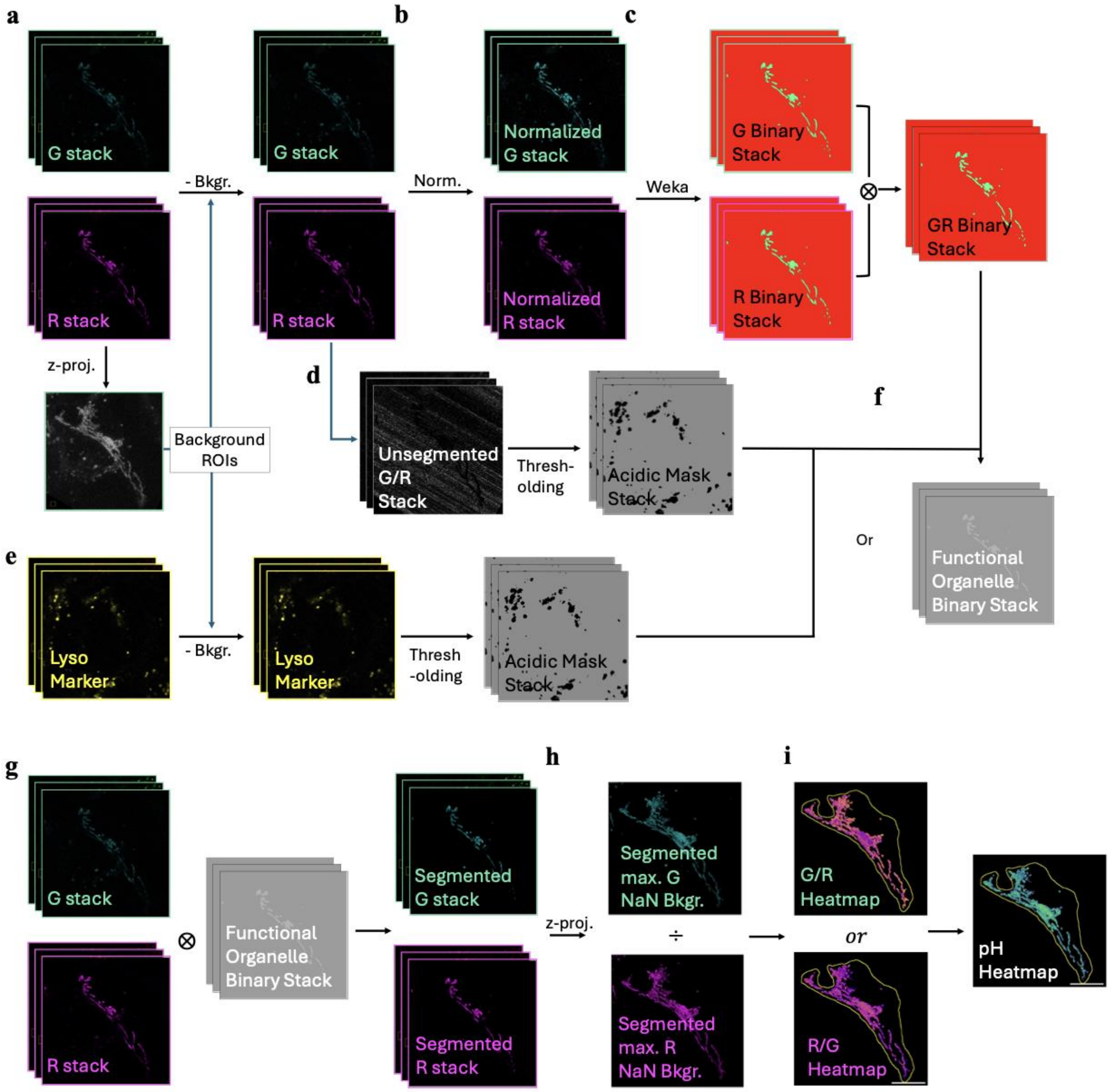


**Extended Data Fig. 2 | SeRapHin image processing pipeline for pH measurement and obtaining pH heatmap. a**, Background subtraction. In max. projection of the 3D stack image, three square ROIs of 30-pixel x 30-pixel size with minimum mean value were identified. The ROI's mean value for each individual plane in the 3D stack image is subtracted as background for each plane. **b**, normalization for segmentation consistency. Mean value from significant pixels selected by auto-thresholding (Otsu method) was set as specific intensity of the image, and the image was normalized so the specific intensity will match the standard intensity (set value for a weka model for each compartment to give best segmentation results) **c**, 3D Weka segmentation done on G and R channel[57]. (Only on R for mitochondria with high R/G, only on G for Golgi with high G/R). For neutral organelles (peroxisome, ER), resulting binary images from G and R are multiplied to yield binary image that sets pixels with both significant G and R as 1 and others 0. For nucleus compartments, 2D weka segmentation was done on z-projected images due to low signal intensity[57]. **d**, For neutral/basic organelles (mitochondrial matrix, peroxisome, and ER), lysosomal signals were identified and removed by thresholding with G/R. 3D binary image stack of pixels with G/R higher than 2.7 (corresponds to pH ~5.5) as 0 and others 1. The clusters of acidic pixels are made into filled patches using Remove Outliers function. **e**, For acidic organelle (Golgi apparatus) and nucleus compartments, lysosomal signals were identified and removed by using DQ-BSA or Dextran-AlexaFluor 647$^{TM}$ (10kDa) as a lysosome marker. The unidirectional bleed-through from lyso-marker channel to SeRapHin (R) was subtracted. Lyso-marker channel thresholding yielded binary image of lysosome pixels made into filled patches of 0, using Remove Outliers function. **f**, Final functional organelle binary stack obtained by multiplying binary stack from weka segmentation and the binary lysosome mask stack. **g**, Background subtracted, unnormalized G and R stack multiplied with functional organelle binary stack to yield segmented G and R stacks. **h**, z-projection and converting background pixels to NaN for calculation of mean G and mean R for the whole segmented organelles in a cell. **i**, G/R or R/G heatmap obtained from mean-filtered, segmented max.G and R images are used to obtain the pH heatmap. G/R (or R/G) conversion to pH demonstrated in **Extended Data Fig. 3**

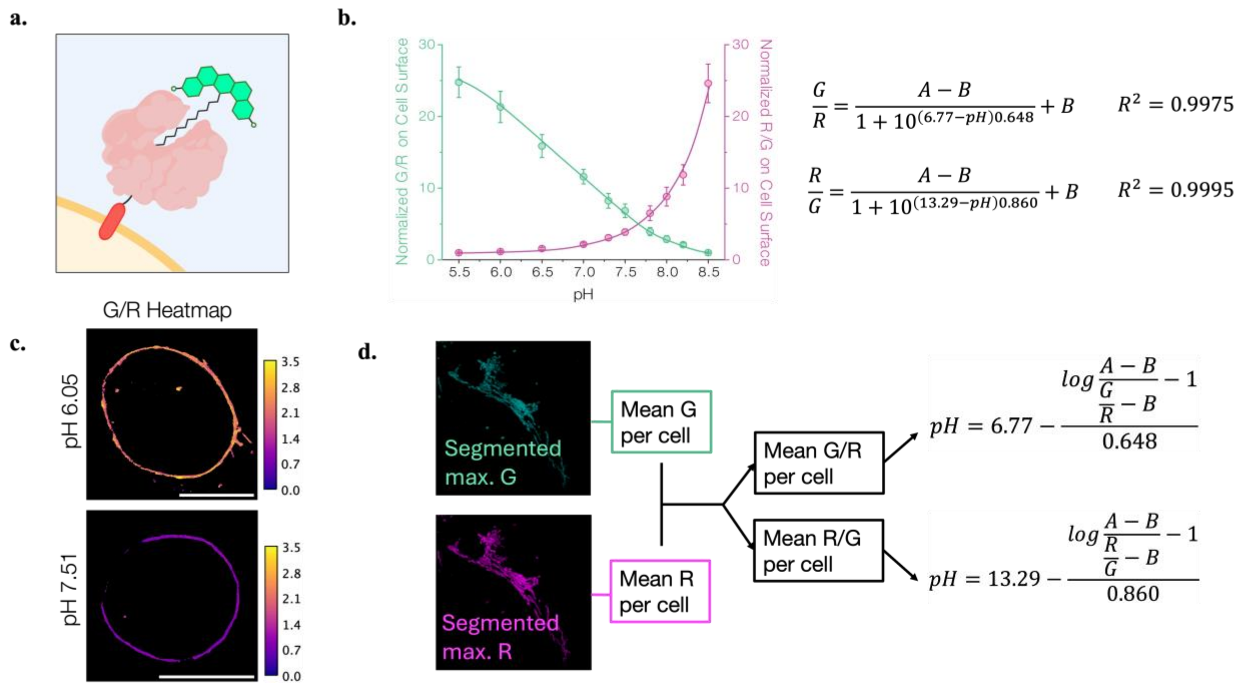


**Extended Data Fig. 3 | pH calculation from SeRapHin G and R value of organelle image. a**, For the cell surface calibration, SeRapHin (green) is localized on the extracellular surface by reacting with HaloTag7 (HT7, salmon) expressed as a chimera with the transmembrane domain of PDGFRβ subunit (red) **b**, G/R - pH and R/G - pH plots fit to sigmoidal function, determining two variables and leaving two variables (A. B) for compensating variance in microscope performance. **c**, A, B are determined by separate experiment (temporally close to the organelle pH imaging) measuring G/R (or R/G) of cell surface immobilized SeRapHin in two known pH points. G/R heatmaps from exemplar pH points are shown. scale bars: 10 μm, **d**, The Mean G and R per cell calculated from Segmented max. projected G and R image from **Extended Data Fig. 2** is used to obtain G/R or R/G. The value is plugged into the pH - G/R function (ER, peroxisome, Golgi and nucleus compartments) or pH – R/G function (mitochondria) to obtain the pH value.

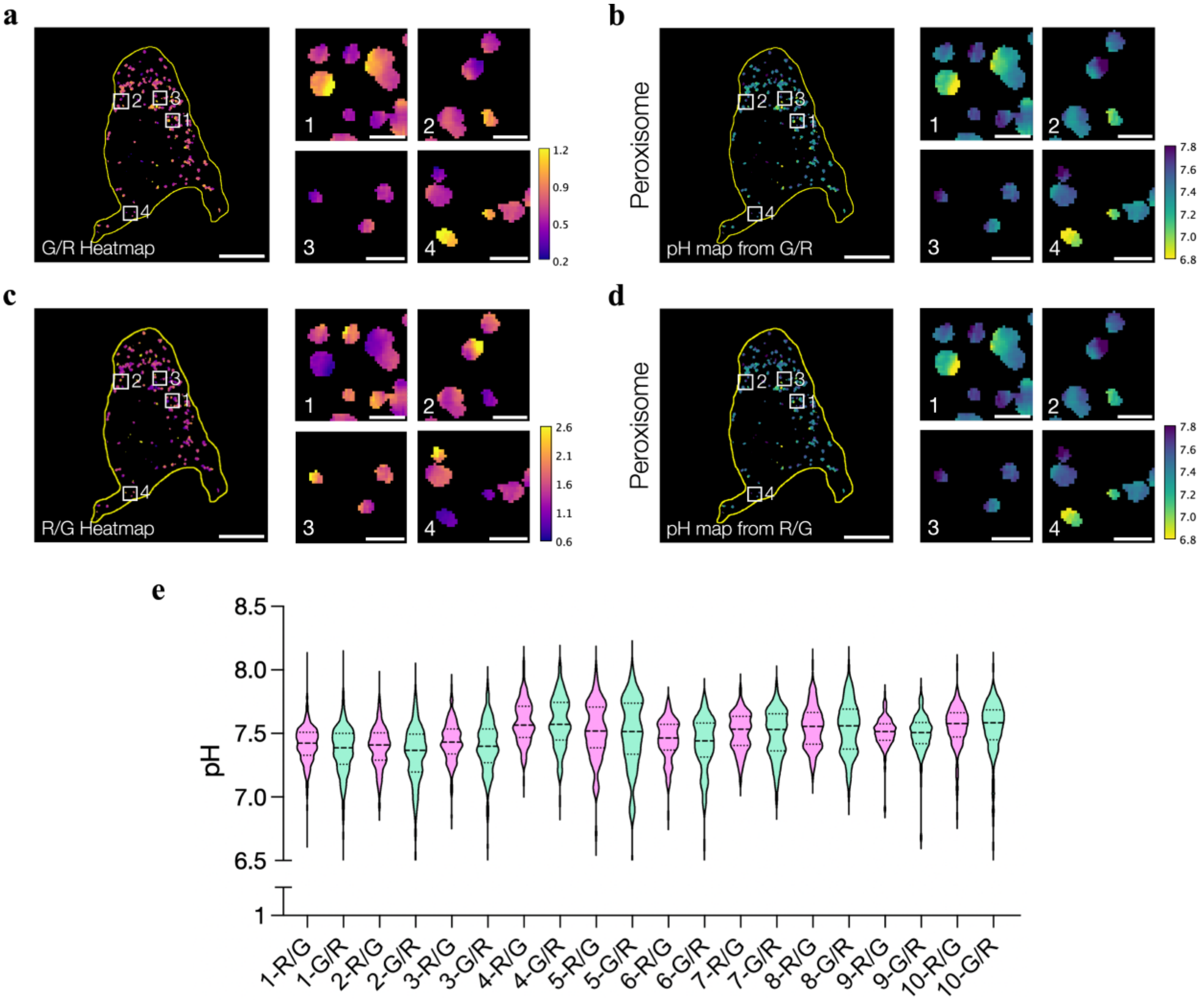


**Extended Data Fig. 4 | Peroxisomal pH measurement is consistent regardless of using G/R or R/G of SeRapHin. a**, G/R Heatmap of peroxisomes of a representative cell. Boxes 1 – 4 are zoomed at the right. **b**, pH heatmap obtained from G/R heatmap in **a.** Boxes 1 – 4 are zoomed at the right. **c,** R/G heatmap of the same cell, showing the opposite pattern from the G/R heatmap. Boxes 1 – 4 are zoomed at the right. **d**, pH heatmap obtained from R/G heatmap show identical pH pattern for each peroxisome. Boxes 1 – 4 are zoomed at the right. All scale bars: 10 μm, all zoom scale bars: 1 μm. Cell outlines are in yellow. **e,** pH values of of individual peroxisomes from ten representative cells calculated using R/G or G/R. For each and every cell, the mean and spread of individual peroxisomal pH values match for both methods using R/G and G/R. (n = 70 – 330 peroxisomes per cell)

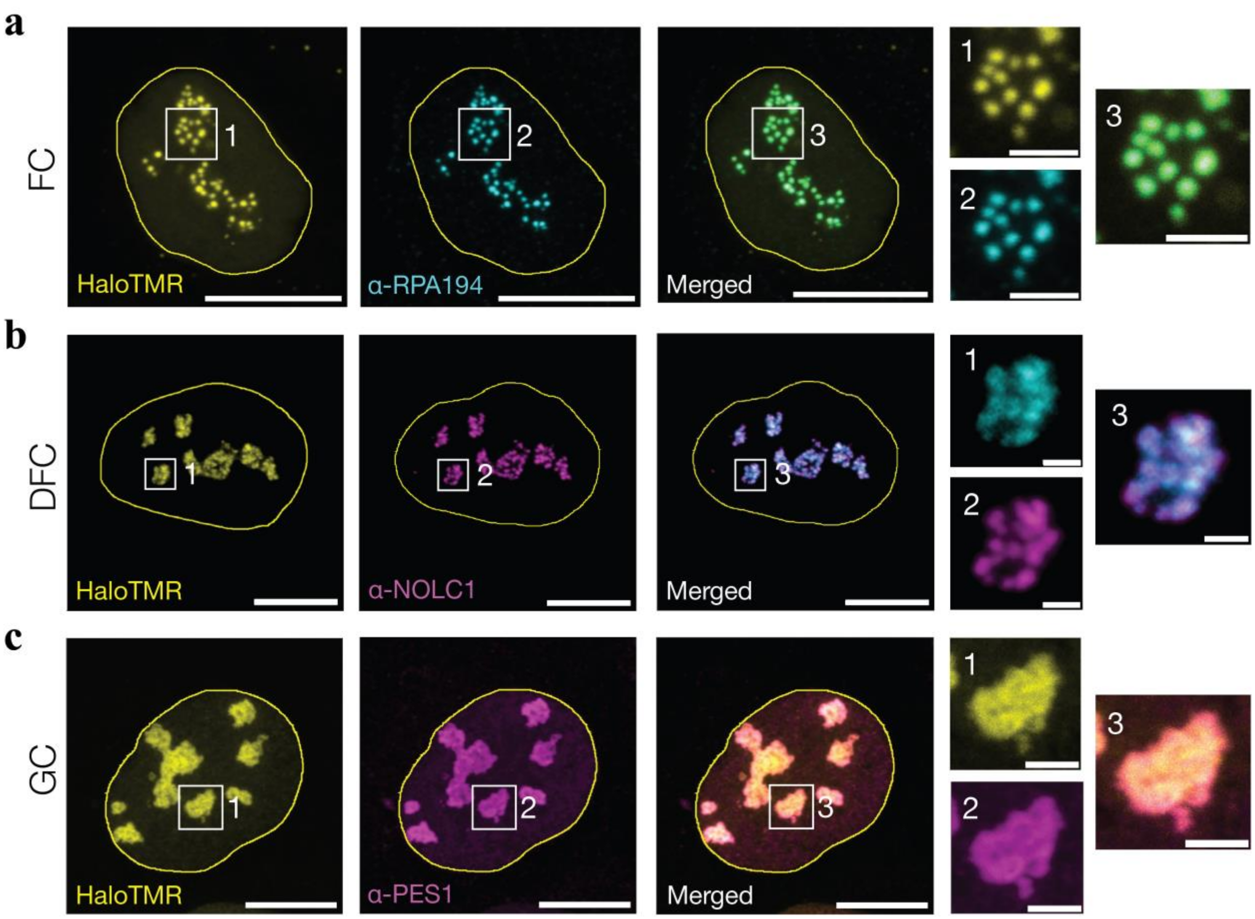


**Extended Data Fig. 5 | HT7 chimeras for sub-nucleolar targeting colocalizes with the markers of corresponding sub-nucleolar regions. a,** Representative images of cells showing colocalization of HT7-RPA43 labeled with HaloTMR (yellow) and the FC marker α-RPA194 (cyan). Merged image is also shown Zoomed images are labeled as 1 – 3. **b**, Representative images of cells showing colocalization of HT7-FBL labeled with HaloTMR (yellow) and the DFC marker α-NOLC1 (magenta). Merged image is also shown Zoomed images are labeled as 1 – 3. **c**, Representative images of cells showing colocalization of NPM1-HT7 labeled with HaloTMR (yellow) and the GC marker α-PES1 (magenta). Merged image is also shown Zoomed images are labeled as 1 – 3. All scale bars = 10 μm. All zoomed in scale bars = 2 μm.
