## Supplementary information for "Organelles harbour pH gradients"

### A chemigenetic pH reporter reveals sub-organellar pH gradients

Supplementary Note 6. SeRapHin fluorescence from different protonation states (for Supplementary Figure S6) ……………………………………………………………………………………………………………...14

Supplementary Scheme 6. Synthesis of SeRapHin-Biotin ……………………………….…………………15

Supplementary Note 7. Synthesis of SeRapHin-Biotin (for SI Scheme 6) ……….,,,,……………………...15

Supplementary Figure S7. Characterization of SeRapHin-Biotin …………………………….…………...16

Supplementary Note 8. Image analysis for cell surface calibration (ROI Selection) ……….……………...17

Supplementary Note 9. Colocalization evaluation (Rotation method) ……………….……………………..17

Supplementary Note 10. Optimized SeRapHin labeling of different organelles ……………………………...18

Supplementary Note 11. Conditions to image Organellar pH …………………………………….……………19

Supplementary Figure S8. Colocalization of B4GalT-HT7 and 58K …………………………….……………..20

Supplementary Note 12. Colocalization of B4GalT-HT7 and 58K (for Supplementary Figure S8) ………...20

Supplementary Note 13. Inter-and intraorganellar pH gradient validation ………….…………………….21

Supplementary Figure S9. Dependence of ln(G/R) of single peroxisomes versus G intensity in live and fixed cells reaffirms pH variation in live peroxisomes …………………………………………………………….22

Supplementary Note 14. Inter-peroxisomal pH variance validation via quantitative analysis ………………22

Supplementary Figure S10. Intra-mitochondrial matrix SeRapHin R/G gradients show continuous patterns over time ……………………………………………………………………………………………………….23

Supplementary Note 15. Continuous SeRapHin R/G patterns over time further validates the intra-mitochondrial matrix pH gradients (for Supplementary Figure S10) ………………………………………..24

Supplementary Note 16. Intra-mitochondrial pH gradient validation via variance quantification ….………..24

Supplementary Note 17. Intra-ER pH gradient validation via variance quantification ………….…………25

**Supplementary Scheme 1. Synthesis of Boc-HaloTag Amine**

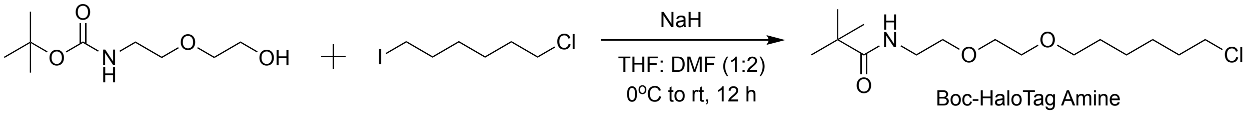

**Supplementary Note 1. Synthesis of Boc-HaloTag Amine (for Supplementary Scheme 1)**

3.5g (1 eq.,17.1 mmol) of 2-[2-(Boc-amino) ethoxy] ethanol was taken in a 100 mL round-bottom flask, and 20 mL of dry THF and 10 mL of dry DMF were added to it. The RB flask was then placed in an ice bath for 30 minutes. Once cooled, 615 mg (1.5 eq., 25.6 mmol) of Sodium hydride was added in parts, and the reaction mixture was stirred for another hour. 3.5 mL (1.4 eq, 23.9 mmol) of 6-chloro-1-iodohexane was then added dropwise, and the reaction was stirred overnight at room temperature. Then the solvent was removed, and the compound was extracted using Ethyl acetate (3 x 150 mL) and water, dried over sodium sulfate. The crude product was then purified on silica gel column chromatography using Hexane: Ethyl acetate (80:20), and the pure Boc-HaloTag amine (2.5g, 45%) was collected. It was then characterized using LRMS and NMR.

$^{1}$H NMR (400 MHz, $CDCl_3$, δ ppm): 4.93 (s, 1H), 3.56 – 3.52 (m, 2H), 3.52 – 3.44 (m, 6H), 3.40 (t, J = 6.7 Hz, 2H), 3.25 (q, J = 5.3 Hz, 2H), 1.75 – 1.67 (m, 2H), 1.60 – 1.50 (m, 2H), 1.38 (s, 13H).

Calculated mass (M + $K^+$): 346.15; Experimental: 346.2

a.

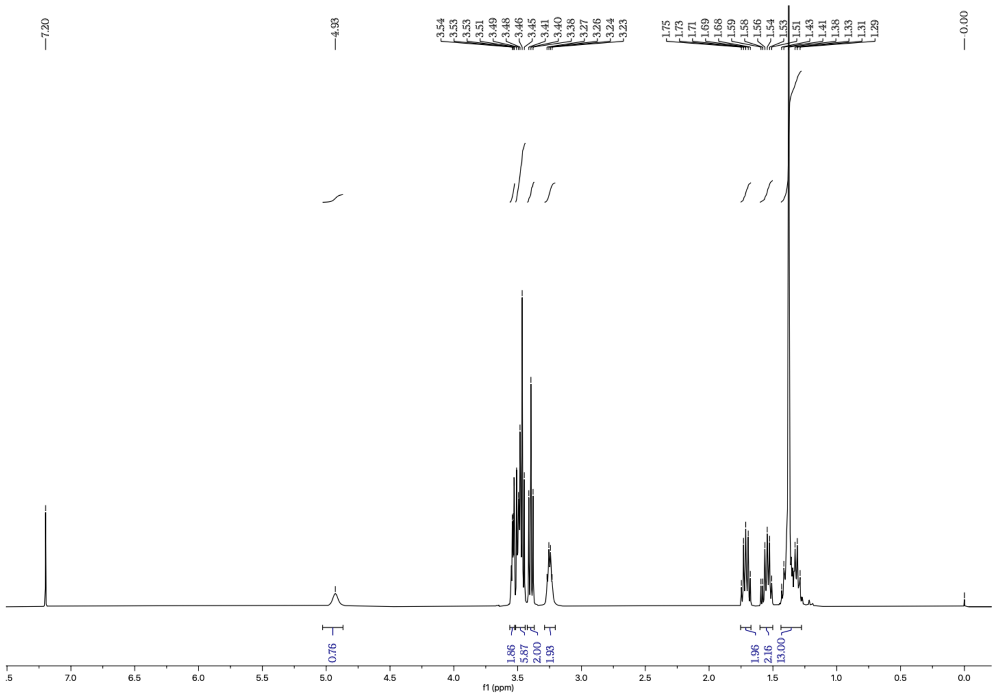

b.

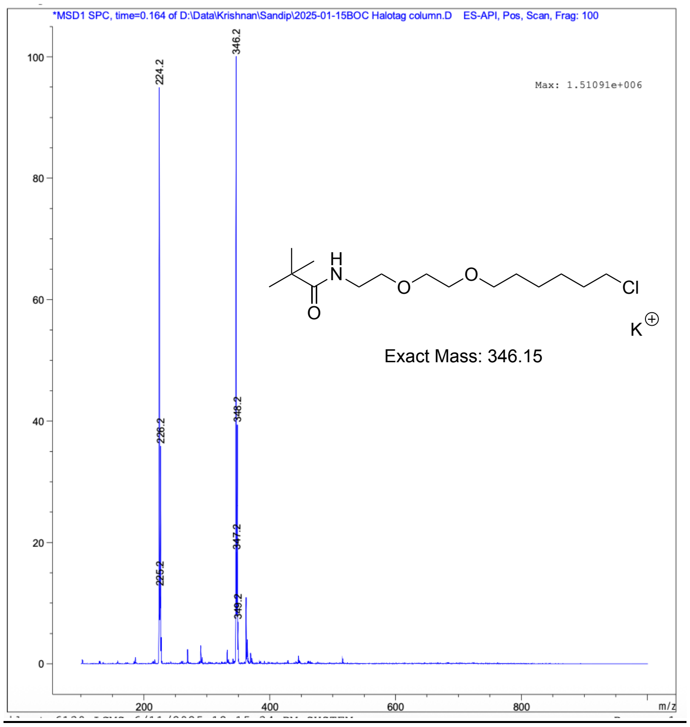

**Supplementary Figure S1. Characterization of Boc-HaloTag Amine a.** $^{1}$H NMR data of Boc-HaloTag Amine **b.** NRMS data of Boc-HaloTag Amine

**Supplementary Scheme 2. Synthesis of HaloTag Amine**

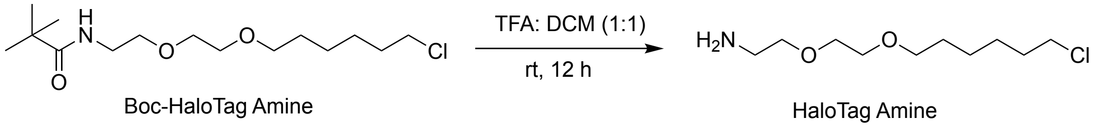

**Supplementary Note 2. Synthesis of HaloTag Amine (for Supplmentary Scheme 2)**

5 mL of DCM was taken in a 50 mL round-bottom flask, and 5 mL of trifluoroacetic acid was added to it. Then, 500 mg (1.6 mmol) of Boc-HaloTag amine was added to it, left with stirring overnight at room temperature. The reaction mixture was then neutralized with sodium hydroxide solution, and the compound was extracted using DCM (3 x 100 mL), dried over sodium sulfate. Solvent was removed to obtain pure HaloTag amine (300 mg, 82%). It was then characterized using LRMS and NMR.

$^1$H NMR (400 MHz, $CDCl_3$, δ ppm): 3.57 – 3.49 (m, 4H), 3.46 (q, $J$ = 6.0 Hz, 4H), 3.40 (t, $J$ = 6.6 Hz, 2H), 2.80 (s, 2H), 1.76 – 1.60 (m, 4H), 1.56 – 1.50 (m, 2H), 1.44 – 1.26 (m, 5H).

Calculated mass (M + $H^+$): 224.14; Experimental: 224.2

a.

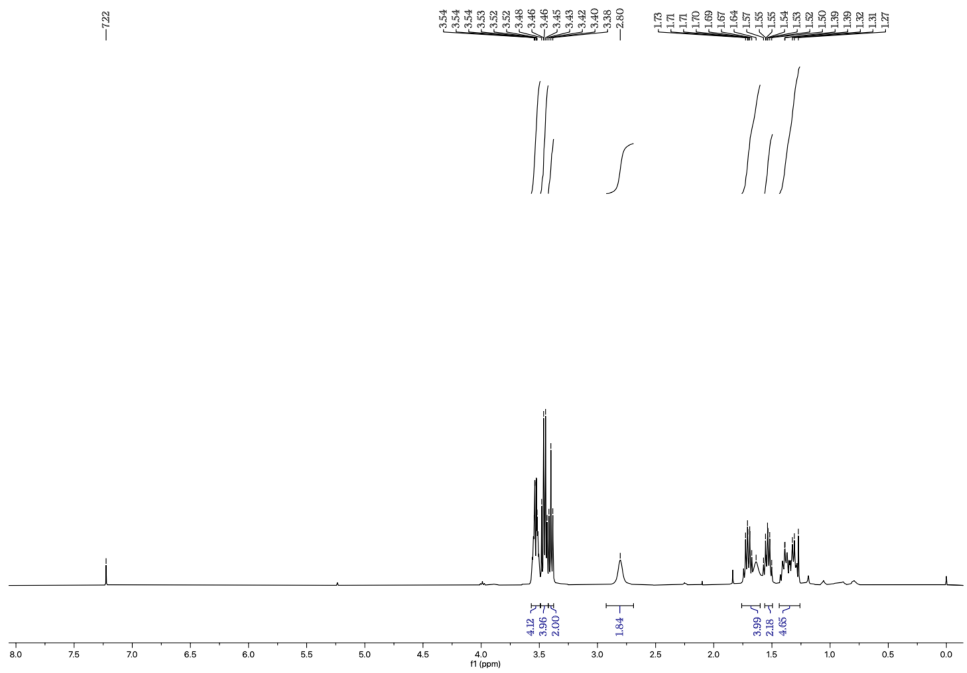

b.

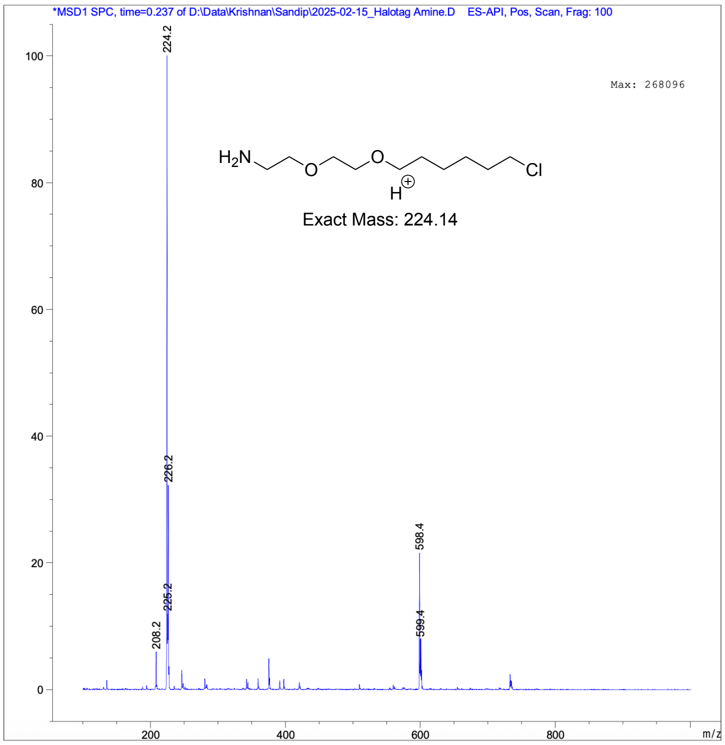

**Supplementary Figure S2. Characterization of HaloTag Amine a**. [1]H NMR data of HaloTag Amine **b.** NRMS data of HaloTag Amine

**Supplementary Scheme 3. Synthesis of PyriOH-$C_6$-COOH**

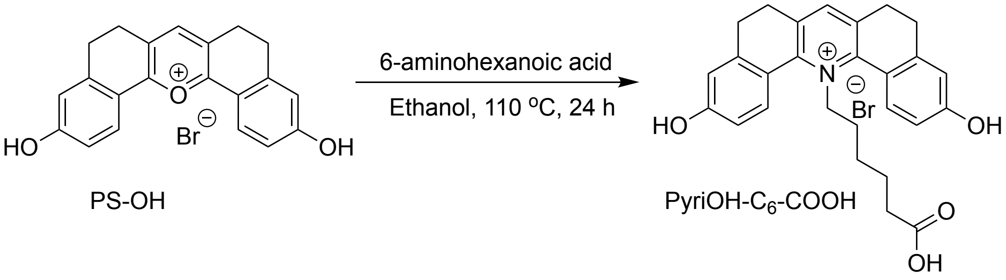

**Supplementary Note 3. Synthesis of PyriOH-$C_6$-COOH (for Supplementary Scheme 3)**
100 mg (1 eq, 0.25 mmol) of PS-OH[1] was taken in a 20 mL pressure tube and dissolved in 3 mL of Ethanol. 22.5 mg (0.9 eq, 0.17 mmol) 6-aminohexanoic acid was added to it. The tube was sealed properly and heated at 90 °C for 12 hours. After that, ethanol was evaporated, and the crude compound was purified on silica gel column chromatography using Dichloromethane: Methanol (92:8), and the pure PyriOH-$C_6$-COOH (50 mg, 39 %) was collected. It was then characterized using LRMS and NMR.

$^1$H NMR (400 MHz, $CDCl_3$, δ ppm): 7.99 (s, 1H), 7.93 (d, $J$ = 8.6 Hz, 2H), 6.88 – 6.80 (m, 4H), 5.10 (t, $J$ = 6.6 Hz, 2H), 2.99 – 2.63 (m, 8H), 1.90 (t, $J$ = 7.2 Hz, 2H), 1.22 (t, $J$ = 7.3 Hz, 2H), 1.09 (t, $J$ = 7.6 Hz, 2H), 0.56 (t, $J$ = 7.7 Hz, 2H).

Calculated mass ($M^+$): 430.2; Experimental: 430.2

a.

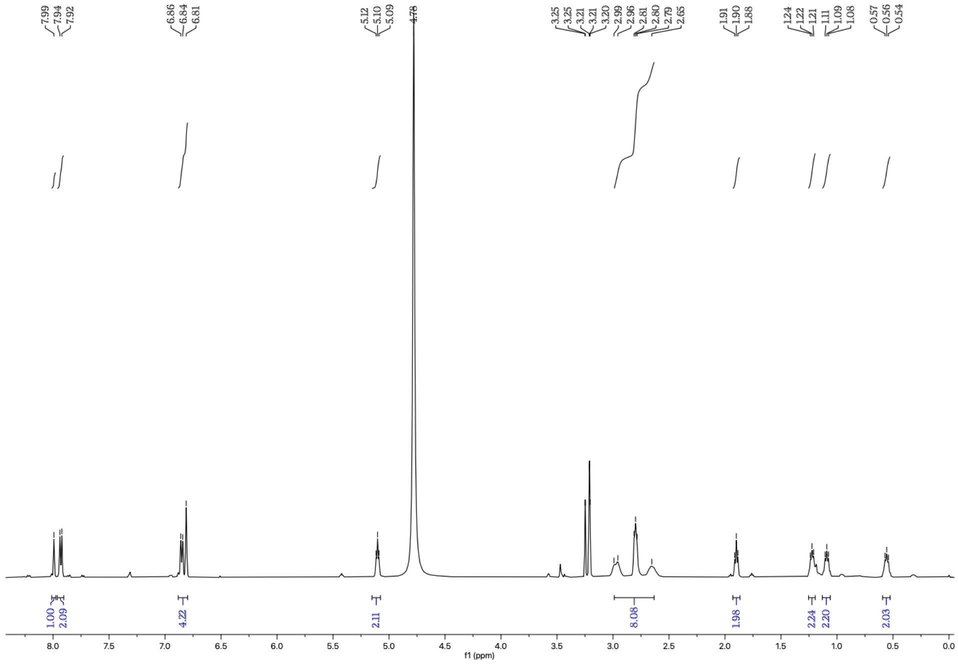

b.

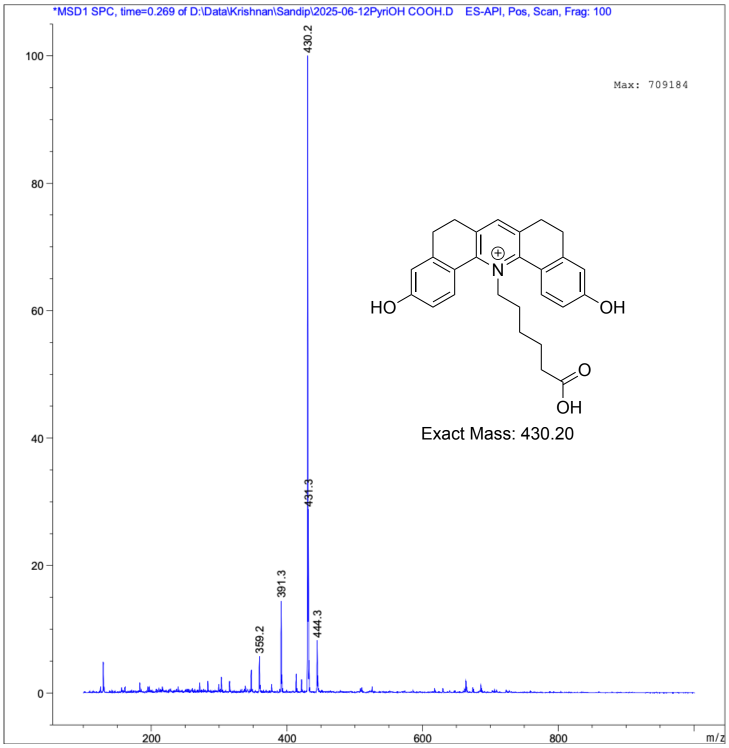

**Supplementary Figure S3. Characterization of PyriOH-C$_6$-COOH a.**[1]H NMR data of PyriOH-C$_6$-COOH **b.** NRMS data of PyriOH-C$_6$-COOH

**Supplementary Scheme 4. Synthesis of PyriOMe-$C_6$-COOH**

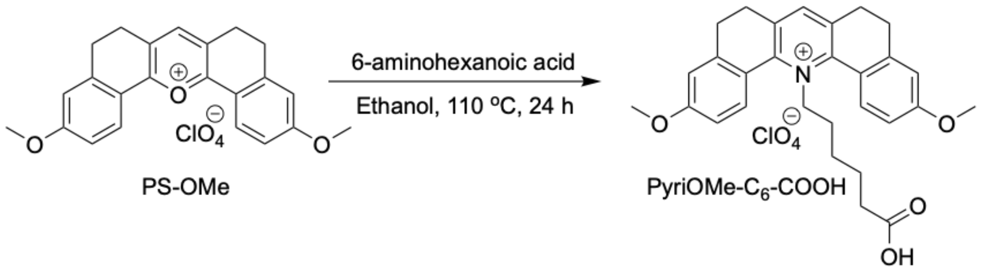

**Supplementary Note 4. Synthesis of PyriOMe-$C_6$-COOH (for Supplementary Scheme 4)**
100 mg (1 eq, 0.22 mmol) of PS-OMe[1] was taken in a 20 mL pressure tube and dissolved in 3 mL of Ethanol. 88.5 mg 6-aminohexanoic acid (3 eq, 0.68 mmol) was added to it. The tube was sealed properly, heated at 90 °C for 12 h. After that, cold Diethyl ether was added to it, and the precipitate was collected using filtration. Then, the precipitate was redissolved using dichloromethane to remove the unreacted 6-aminohexanoic acid. Dichloromethane was evaporated to obtain pure PyriOMe-$C_6$-COOH (93 mg, 74 %). It was then characterized using LRMS and NMR.

$^1$H NMR (400 MHz, $CDCl_3$, δ ppm): 8.07 – 7.95 (m, 3H), 7.03 (dd, *J* = 8.8, 2.7 Hz, 2H), 6.86 (d, *J* = 2.6 Hz, 2H), 5.14 (t, *J* = 6.6 Hz, 2H), 3.86 (s, 6H), 3.14 – 2.62 (m, 8H), 2.04 (t, *J* = 6.9 Hz, 2H), 1.19 (d, *J* = 3.6 Hz, 2H), 0.84 – 0.73 (m, 2H), 0.60 (tt, *J* = 10.1, 6.4 Hz, 2H).

Calculated mass ($M^+$): 458.23; Experimental: 458.3

a.

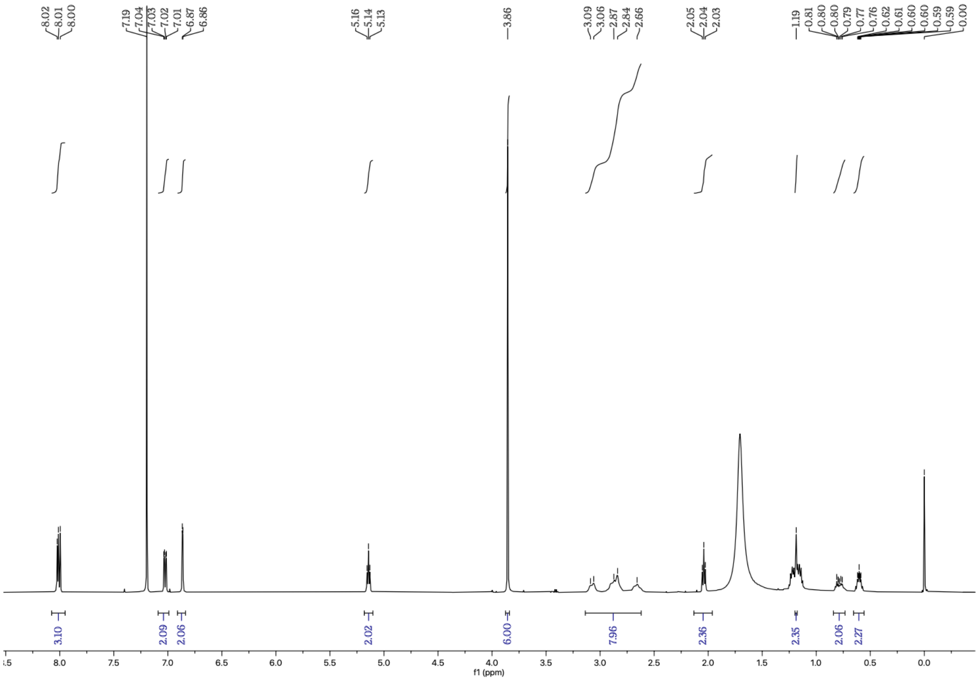

b.

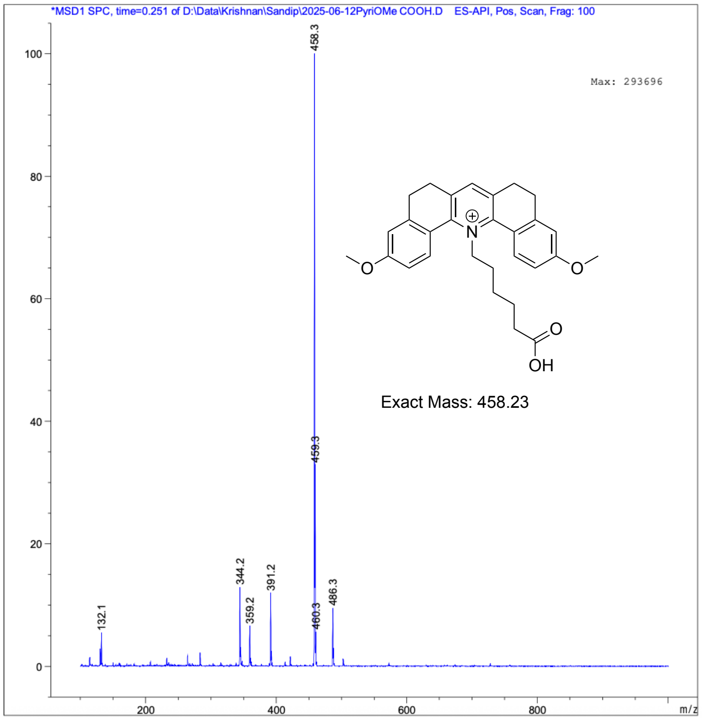

**Supplementary Figure S4. Characterizationof PyriOMe-C₆-COOH a.** [1]H NMR data of PyriOMe-C₆-COOH **b.** NRMS data of PyriOMe-C₆-COOH

**Supplementary Scheme 5. Synthesis of SeRapHin**

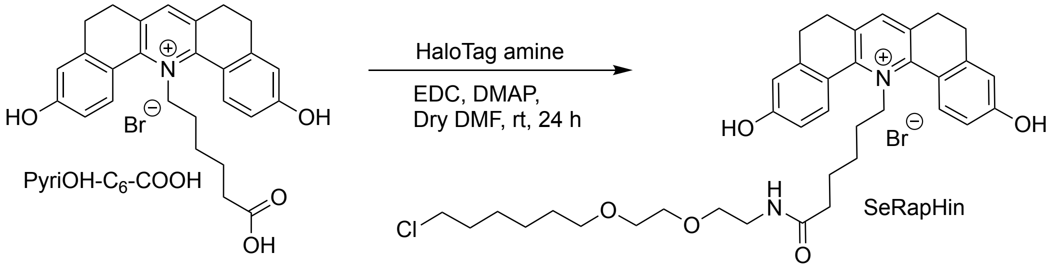

**Supplementary Note 5. Synthesis of SeRapHin (for Supplementary Scheme 5)**

20 mg of PyriOH-$C_6$-COOH (1 eq, 0.04 mmol) was taken in a 25 mL round-bottom flask and dissolved in 5 mL of dry DMF. 12.15 mg of EDC (2 eq., 0.08 mmol) and 9.56 mg of DMAP (2 eq, 0.08 mmol) were added to it. The reaction mixture was stirred on an ice bath for 30 minutes. Then, 50 mg of HaloTag amine (5.7 eq., 0.09 mmol) was added to it and kept with stirring for 24 h. Once the completion of the reaction was confirmed by checking TLC, DMF was evaporated, and the crude compound was purified on silica gel column chromatography using Dichloromethane: Methanol (96:4), and the pure SeRapHin (12 mg, 42%) was collected. It was then characterized using LRMS and NMR.

$^{1}$H NMR (400 MHz, MeOD, δ ppm): 8.00 (s, 1H), 7.92 (d, *J* = 8.6 Hz, 2H), 6.89 – 6.79 (m, 4H), 5.09 (t, *J* = 6.6 Hz, 2H), 3.41 – 3.34 (m, 4H), 3.24 – 3.14 (m, 8H), 2.99 – 2.65 (m, 8H), 1.83 (t, *J* = 7.2 Hz, 2H), 1.70 – 1.60 (m, 2H), 1.48 (p, *J* = 6.7 Hz, 2H), 1.37 – 1.20 (m, 6H), 1.12 (q, *J* = 7.5 Hz, 2H), 0.55 (dd, *J* = 8.8, 6.2 Hz, 2H).

$^{13}$C NMR (101 MHz, MeOD, δ ppm): 174.0, 161.7, 153.2, 144.2, 141.2, 136.2, 130.4, 119.8, 115.0, 114.8, 70.8, 69.8, 69.8, 69.1, 44.3, 38.9, 34.8, 32.3, 29.1, 28.7, 28.1, 27.6, 26.3, 25.1, 24.6, 24.5.

Calculated mass ($M^+$): 635.32; Experimental: 635.4

a.

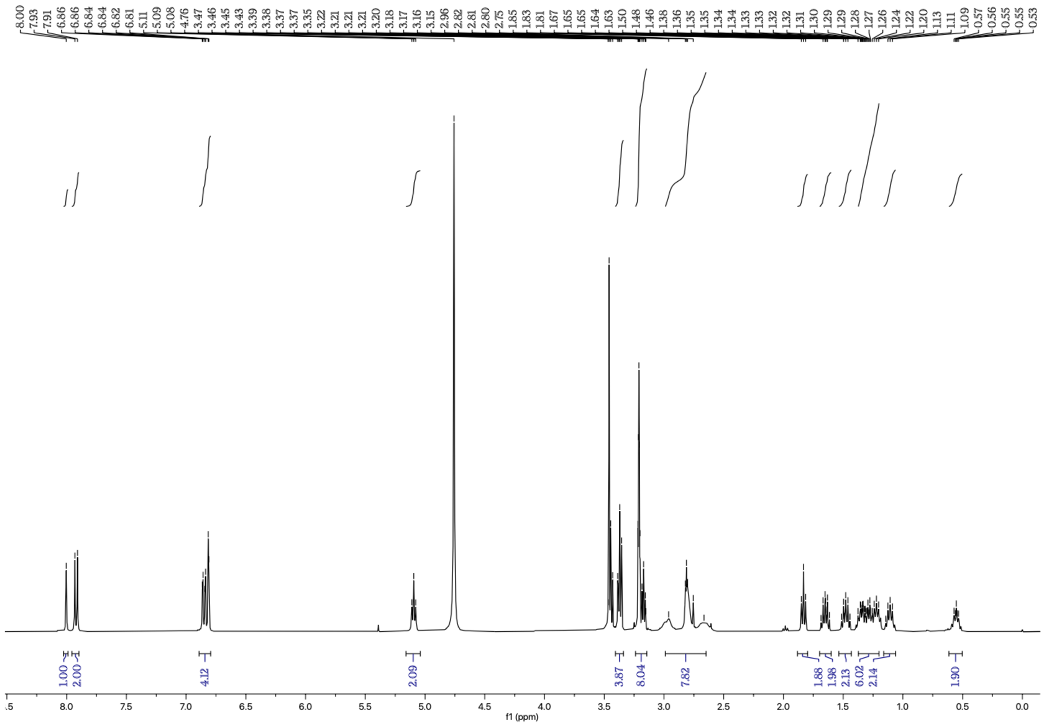
8.00 7.93 7.91 6.86 6.86 6.84 6.84 6.82 6.81 5.11 5.09 5.08 4.76 3.47 3.46 3.45 3.43 3.39 3.38 3.37 3.37 3.35 3.22 3.21 3.21 3.21 3.20 3.18 3.17 3.16 3.15 2.96 2.82 2.81 2.80 2.75 1.85 1.83 1.81 1.67 1.65 1.65 1.64 1.63 1.50 1.48 1.46 1.38 1.36 1.35 1.35 1.34 1.34 1.33 1.33 1.32 1.32 1.31 1.30 1.29 1.29 1.28 1.27 1.26 1.24 1.24 1.22 1.20 1.13 1.11 1.09 0.57 0.56 0.55 0.55 0.53
1.00 2.00 4.12 2.09 3.87 8.04 7.82 1.88 1.98 2.13 6.02 2.14 1.90
f1 (ppm)

b.

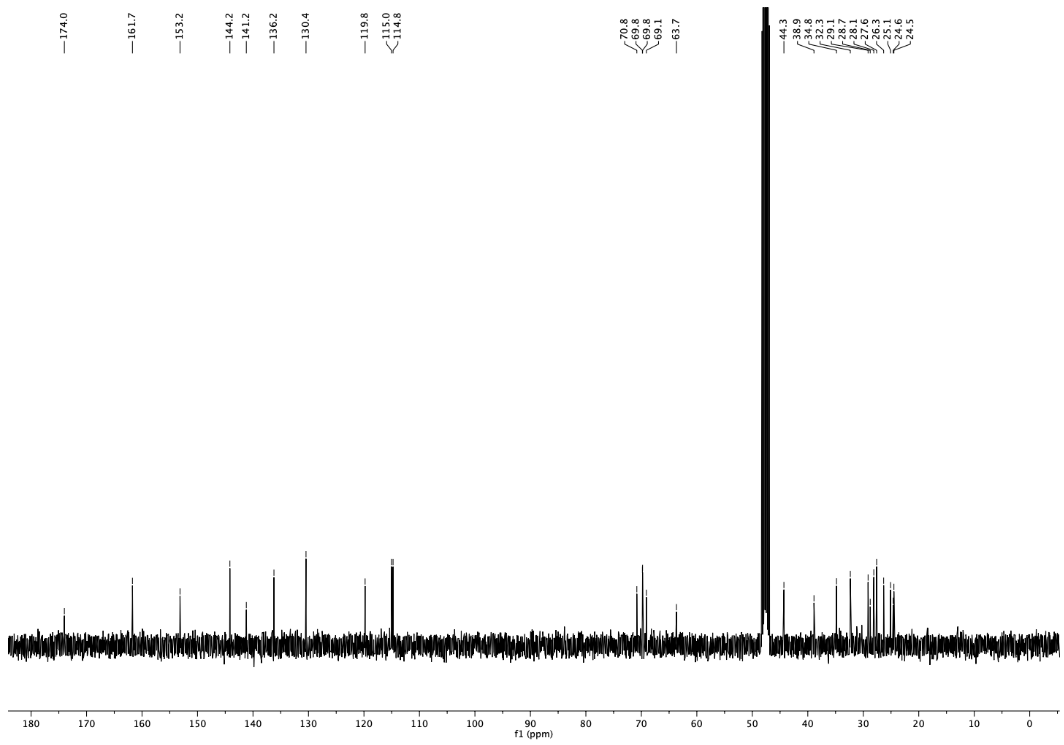
174.0 161.7 153.2 144.2 141.2 136.2 130.4 119.8 115.0 114.8 70.8 69.8 69.8 69.1 63.7 44.3 38.9 34.8 32.3 29.1 28.7 28.1 27.6 26.3 25.1 24.6 24.5
f1 (ppm)

c.

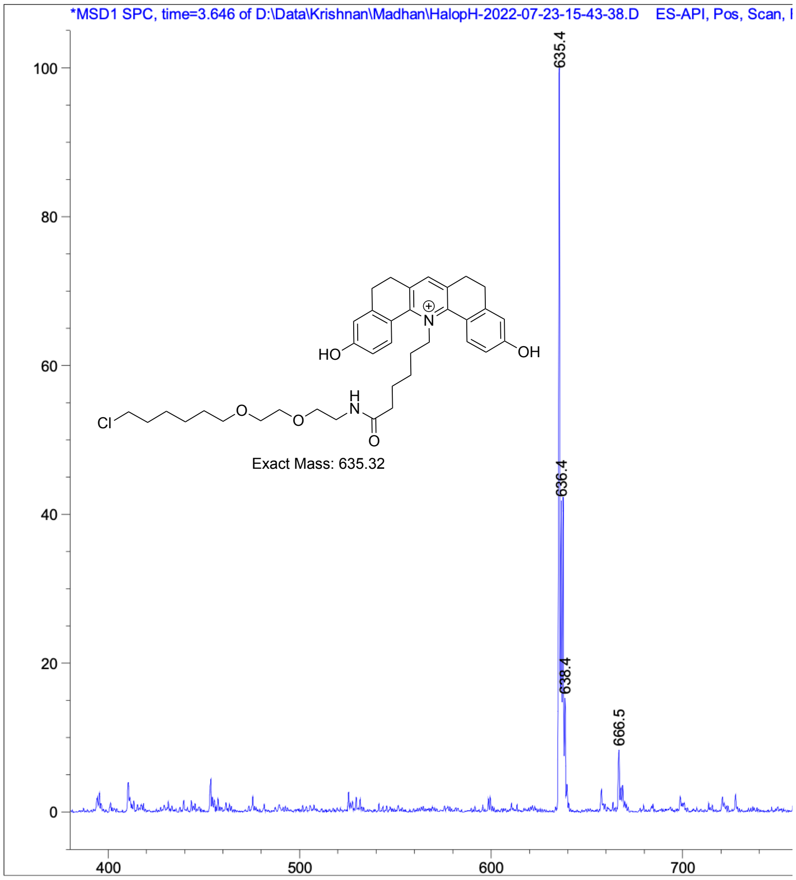

**Supplementary Figure S5. Characterization of SeRapHin a**. [1]H NMR data of SeRapHin **b**. [13]C NMR data of SeRapHin. **c.** NRMS data of SeRapHin

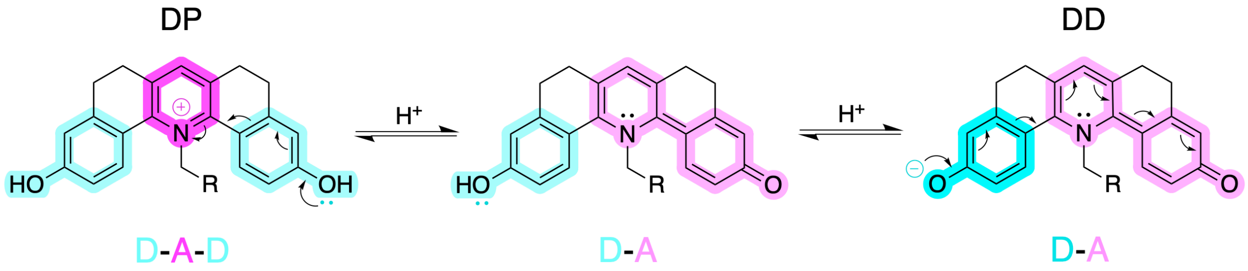

**Supplementary Figure S6. Protonated states of SeRapHin showing donor (D) and acceptor (A) moieties highlighted in cyan and magenta respectively. Left:** Doubly protonated state (DP) is a Donor-Acceptor-Donor (D-A-D) system with green fluorescence. Centre: Mono-protonated state is a Donor-Acceptor (D-A) system. **Right:** Doubly deprotonated (DD) state is also Donor-Acceptor (D-A) system with red fluorescence, but the DD and DP acceptors differ. Higher donor/acceptor strength is indicated by a more intense color.

**Supplementary Note 6. SeRapHin fluorescence from different protonation states (for Figure S6)**

In the doubly protonated (DP) state, SeRapHin shows a strong green fluorescence ($\lambda_{Em}$ = 540 nm) due to the intramolecular charge transfer (ICT) from the two symmetric donor hydroxyl groups adjacent to the acceptor pyridinium ring in the center yielding a D-A-D system. When the first deprotonation ($pKa_1$) happens, the symmetry of SeRapHin breaks, and the quinone formed by the tautomerization of phenoxide becomes the new acceptor. When the second deprotonation ($pKa_2$) happens at a higher $pKa_2$ it gives the doubly deprotonated (DD) state whose emission is red shifted ($\lambda_{Em}$ = 625 nm) because the phenoxide in DD is a stronger donor compared to the hydroxyl group in DP. The new acceptor is now the carbonyl oxygen.

The two non-aromatic rings in DP act as a bridge to facilitate efficient electron donation from the hydroxyl donors to the pyridinium acceptor. With the first deprotonation, three rings become conjugated, and quinone becomes the new acceptor. When SeRapHin is in the DD state, the structural uniqueness restricts it from becoming a completely conjugated system. In DD, the two conjugated rings work as an acceptor, while the phenoxide works as a strong donor, and the remaining non-aromatic rings bridge the new D-A system.

**Supplementary Scheme 6. Synthesis of SeRapHin-Biotin**

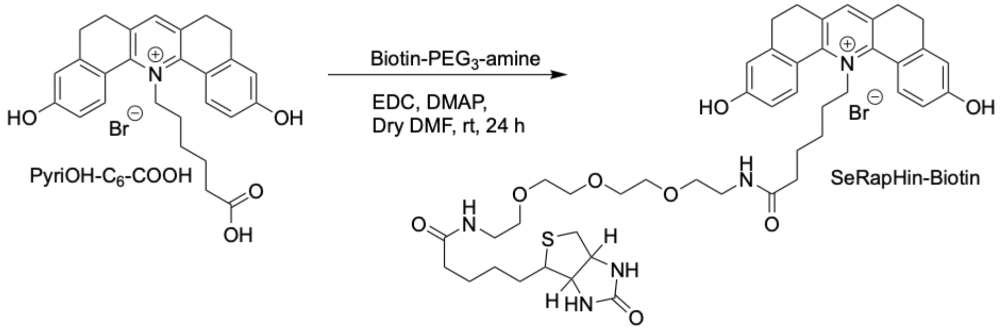

**Supplementary Note 7. Synthesis of SeRapHin-Biotin (for Supplementary Scheme 6)**

10 mg of PyriOH-$C_6$-COOH (1 eq, 0.02 mmol) was taken in a 25 mL round-bottom flask and dissolved in 5 mL of dry DMF. 6 mg of EDC (2 eq, 0.04 mmol) and 5 mg of DMAP (2 eq, 0.04 mmol) were added to it. The reaction mixture was stirred on an ice bath for 30 minutes. Then, 15 mg of Biotin-$PEG_3$-amine (1.8 eq., 0.04 mmol) was added to it and kept with stirring for 24 h. Once the completion of the reaction was confirmed by checking TLC, DMF was evaporated, and the crude compound was purified on silica gel column chromatography using Dichloromethane: Methanol (93:7), and the pure SeRapHin Biotin (4 mg, 22%) was collected. It was then characterized using LRMS and NMR.

Calculated mass ($M^+$): 830.42; Experimental: 830.4

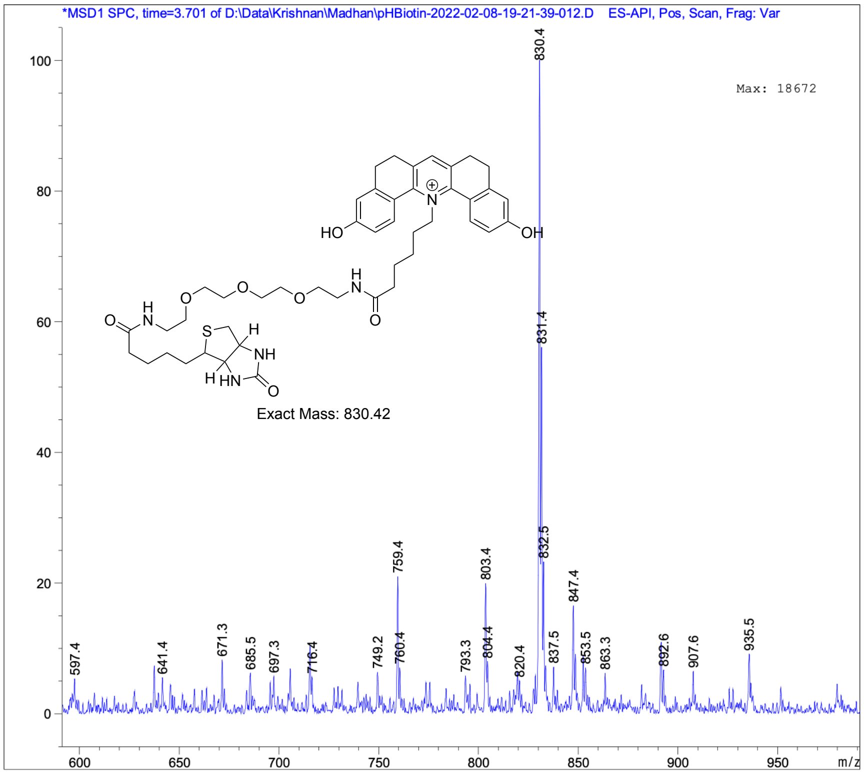

**Supplementary Figure S7. Characterization of SeRapHin-Biotin** NRMS data of SeRapHin-Biotin.

**Supplementary Note 8. Image analysis for cell surface calibration (ROI selection)**

HT7 with IgK signal sequence and PDGFRβ transmembrane sequence localizes mostly in the plasma membrane. However, a middling population is present in secretory organelles due to the process of membrane protein biogenesis. Further, due to constitutive endocytosis, a small population is also present in endosomes. Hence SeRapHin signal is also observed from these regions inside the cell. Hence it is critical to select ROIs exclusively from the signal at the plasma membrane for calibration corresponding to the extracellular buffer. Weka segmentation was used to create binary image, and 4 – 6 regions of plasma membrane that does not include apparent intracellular acidic vesicles were selected as ROIs for G/R or R/G calculations.

**Supplementary Note 9. Colocalization evaluation (Rotation method)**

To evaluate the colocalization, we calculated the Pearson's correlation coefficient (PCC) on the 3D image stack using drawn cell ROIs. To test if the PCC value is coming from random colocalization, we performed pixel shift (15 pixels) on one of the channels and re-calculated the PCC. The pixel shift method is suitable for punctate organelles like peroxisomes since signals in two channels from such objects do not overlap after a pixel shift. However, this approach fails for mitochondria, ER and the Golgi apparatus since signal overlap does not change drastically by pixel shift due to their long or wide morphology. To test for the random colocalization, we instead applied a rotation method where we rotate one of the channels by $90^{o}$. We cannot directly use the drawn cell ROI since the included pixels in the ROI before and after the rotation can be quite different. To solve this, we make a new circular ROI that uses the center of mass of the cell ROI as its center and has the same area with the cell ROI. PCC was calculated using the circular ROI before and after $90^{o}$ rotation to evaluate colocalization and cross-verify whether it was due to random colocalization. The $90^{o}$ rotation was performed using the circular ROI's center as the rotation center.

**Supplementary Note 10. Optimized SeRapHin labeling of different organelles**

*Mitochondria, ER, and peroxisome*

HeLa cells seeded on 0.8 $cm^2$ area of glass-bottomed imaging dish to ~70% confluency were transfected with 250 ng of the HaloTag plasmid for the organelle of interest using Lipofectamine3000 following the manufacturer's protocol. After 18 – 24 h from transfection, the imaging dish was washed 3 times with 1mL PBS each time. The cells were then treated with 1 μM SeRapHin in serum-free DMEM for 2 h in 37°C, 5% $CO_2$. It is then washed 3 times with 1mL PBS each time and incubated in complete media (DMEM without phenol red with 10% FBS) in 37°C, 5% $CO_2$ for over 12 h.

*Golgi apparatus*

To minimize leaky expression and specifically localize HT7 in the target Golgi sub-compartment, we must lower protein expression yet retain adequate SeRapHin signal to noise ratio. This was achieved by reducing the amount of the HT7-fusion plasmid and complementing it with an empty "filler" vector to decrease of the HT7 incorporation per plasmid-liposome complex.

With HaloTag technology one labels that specific population of HT7-protein that is expressed during incubation with SeRapHin. Thus, there is a tight time-window to wash out free, unreacted SeRapHin, otherwise at the time of imaging, a significant portion of the labeled B4GalT-HT7 will have trafficked onwards and out of the TGN. We therefore shortened washing times to 2 h to remove the signal from unreacted SeRapHin in some locations e.g., mitochondria, but not others e.g., lysosomes. Importantly, TGN components such as B4GalT-HT7 can also end up in lysosomes due to autophagy, since the TGN is a major source for autophagosome membranes[1]. Because Seraphin-labeling needs serum starvation, which generally elevates autophagy, we limited labeling times for the TGN to 1 h to minimize autophagy. Hence, we used DQ-BSA or Alexa647-Dextran as a mask for lysosomes during image analysis.

HeLa cells on 0.8 $cm^2$ area of glass-bottomed imaging dish to ~70% confluency were transfected with a mixture of 100 ng of either Stx5-HT7, MGAT2-HT7 or B4GalT-HT7 and 150 ng empty vector using Lipofectamine3000 following the manufacturer's protocol. After 18 – 24 h in phenol red-free complete media in 37°C, 5% $CO_2$, the imaging dish was washed 3 times with 1 mL PBS. The cells were then treated with 1 μM SeRapHin and 2 μM DQ-BSA (cis-Golgi and TGN) in serum-free DMEM for 1.5 h in 37°C, 5% $CO_2$. It is then washed 3 times with 1 mL PBS, and incubated in complete media (DMEM without phenol red with 10% FBS) in 37°C, 5% $CO_2$ for 2.5 h. For medial-Golgi, cells were labeled with Alexa647-Dextran for 16 h in HBSS, treated with SeRapHin for 1 h, and then imaged.

*Nucleoplasm and chromatin*

U2OS cells on 0.8 $cm^2$ area of glass-bottomed imaging dish to ~70% confluency were transfected with the mixture of 250 ng of HT7-SKL or HT7-H2B using Lipofectamine3000 following the manufacturer's protocol. After 18 – 24 h in phenol red-free complete media in 37°C, 5% $CO_2$, the imaging dish was washed 3 times with 1mL PBS each time. The cells were then treated with 2 μM SeRapHin and 2 μM DQ-BSA in serum-free DMEM for 1.5 h in 37°C, 5% $CO_2$. It is then washed 3 times with 1mL PBS each time, and incubated in complete media (DMEM without phenol red with 10% FBS) in 37°C, 5% $CO_2$ for 2.5 h.

*Fibrillar (FC), dense fibrillar (DFC) and granular (GC) components of nucleolus*

The FC was labeled using HT7-RPA43 and the procedure described for the nucleus above. For the GC and the DFC, U2OS cells stably expressing either HT7-NPM1 or HT7-FBL were seeded on 0.8 $cm^2$ area of glass-bottomed imaging dish to ~50% confluency. The imaging dish was washed 3 times with 1 mL PBS. The cells were then treated with 2 μM SeRapHin and 2 μM DQ-BSA in serum-free DMEM for 1.5 h in 37°C, 5% $CO_2$. It is then washed 3 times with 1 mL PBS, and incubated in complete media (10% FBS in DMEM, no phenol red) in 37°C, 5% $CO_2$ for 2.5 h.

**Supplementary Note 11. Conditions to image organellar pH**

ER, peroxisome, and Golgi: L-15 media without phenol red was selected for the pH stability in the $CO_2$-free imaging environment. Especially for Golgi, glutamine must be present in the imaging media because it is required to retrieve mis-localized proteins in endosomes to the TGN [2]).

Mitochondria: HBSS was selected as imaging media instead of L-15 to prevent the effects of glucose starvation on mitochondrial activity since L-15 contains galactose instead of glucose. Imaging was done within 1.5 h to minimize the effect of serum starvation.

Nucleus and nucleolus: Since nucleolus can be highly sensitive to serum starvation and temperature change, it is critical to minimize these conditions during SeRapHin treatment and imaging. 10% FBS in DMEM with 20 mM HEPES without phenol red was used as imaging media and imaging was done under 37°C, 5% $CO_2$ to avoid any effect of starvation, heat/cold shock, or pH fluctuations of the extracellular media during imaging. The imaging media was pH stabilized by incubation in 37°C, 5% $CO_2$ for 1 h before addition into the imaging dish. After the imaging dish was transferred into the pre-equilibrated imaging chamber in 37°C, 5% $CO_2$, it was left to stabilize further for 30 mins before images were acquired.

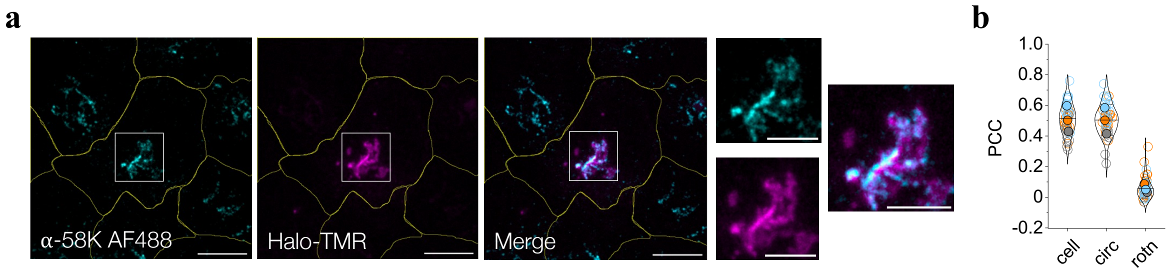

**Supplementary Figure S8. Colocalization of B4GalT-HT7 and 58K a.** Left: Fluorescence images of a HeLa cell transfected wit B4GalT-HT7, labeled with HaloTMR, and immunolabeled with α-58K and Alexa Fluor 488™ conjugated secondary antibody. Right: Smaller panels are zoomed images of the indicated white boxes. Scale bar = 10 μm, zoomed scale bar = 5 μm **b.** Pearson's Correlation Coefficient (PCC) values and upon rotation as described in **Supplementary Note 9**. Open circles are PCC datapoints from a cell using r.o.i.s corresponding to cell outline (cell), circular outline (circ), and upon rotating one channel in the circular outline (rotn). Experiments were performed in triplicates, and each trial is distinguished by color. Filled circles are the color-coded mean value corresponding to data from a given trial.

**Supplementary Note 12. Colocalization of B4GalT-HT7 and 58K (for SI Figure S8)**
To cross-verify localization of B4GalT-HT7 in the Golgi apparatus, we checked its colocalization by immunofluorescence with the endogenous Golgi protein denoted 58K. HeLa cells are transfected with B4GalT-HT7 as described in **Supplementary Note 9** and labeled with Halo-TMR 18 h after transfection following the vendor's protocol. The cells are then fixed with 4% PFA and permeabilized with 0.25% TritonX-100 + 0.1% Tween-20. Cells are immunolabeled with α-58K (abcam, ab27043) and Alexa Fluor 488™ goat-anti-mouse IgG H&L (Invitrogen, A11001). The colocalization of both channels is evaluated by their PCC and tested for random colocalization using the rotation method described in **Supplementary Note 9**. B4GalT-HT7 labeled with HaloTMR showed robust colocalization with α-58K with PCC values from cell r.o.i.s or circular r.o.i.s higher than upon rotating one channel of the circular r.o.i., effectively ruling out random colocalization of B4GalT-HT7 and α-58K.

**Supplementary Note 13. Inter- and intraorganellar pH gradient validation**

To determine whether the observed heterogeneity in organellar pH reflects real differences or are simply the result of an imaging artifact, we collapsed organellar pH either by fixation or V-ATPase inhibition by Bafilomycin treatment. Under either of these conditions, the G/R or R/G values are expected to become uniform throughout the organelle. If imaging artifacts were responsible, variations in G/R. or R/G values would persist even after clamping or fixation. In contrast, the disappearance of variability under these conditions would indicate that any variation observed in live cells represents genuine pH differences.

Peroxisomes: To stop biological processes and collapse pH variation if any between peroxisomes in a cell, we used methanol fixation and permeabilization. HeLa cells are transfected with HT7-SKL and treated with SeRapHin as described in **Supplementary Note 10**. Then the cells are fixed with -20°C methanol for 10 mins. The cells were washed 3 times with 1 mL PBS and incubated in room temperature PBS for 1h before imaging.

Mitochondria: Methanol fixation was also used for collapsing the mitochondrial pH gradient, if any. HeLa cells are transfected with mito-HT7 and treated with SeRapHin as described in **Supplementary Note 10.** Then, the cells are fixed with -20°C methanol for 10 mins. The cells were washed 3 times with 1 mL PBS and incubated in room temperature PBS for 1h before imaging.

Endoplasmic Reticulum: To test if the observed G/R variation in ER depends on V-ATPase activity, we inhibited V-ATPase using Bafilomycin. HeLa cells are transfected with ER-HT7 or and treated with SeRapHin as described in **Supplementary Note 10**. Additionally, the cells were treated with AlexaFluor 647 Dextran (10kDa) during the >12h SeRapHin wash step for masking all endo-lysosomal compartments to avoid any variance arising from insufficiently excluded signal from the endo-lysosomal compartments. Then the cells are treated with 500nM bafilomycin A1 for 1h. The cells were washed 3 times with 1 mL PBS and imaged in L15 media.

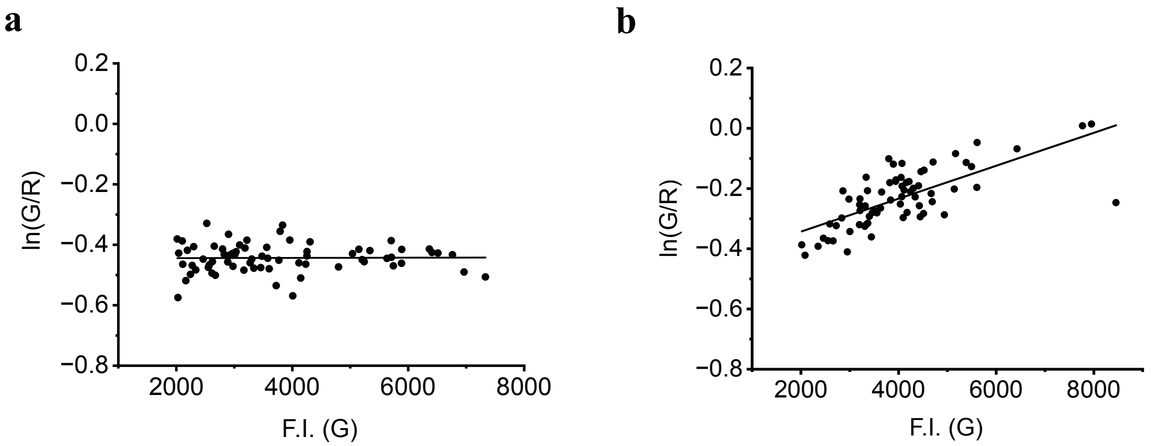

**Supplementary Figure S9. Dependence of ln(G/R) of single peroxisomes versus G intensity in live and fixed cells reaffirms pH variation in live peroxisomes. a.** The ln(G/R) value of single peroxisomes is independent of the intensity in the G channel in a fixed cell and is consistent with Extended Data Fig. 3. **b**, In live cells ln(G/R) value shows an apparent dependence on the G intensity of SeRapHin. All datapoints (peroxisomes) are from a representative fixed or live cells, as applicable with std(ln(G/R)) close to the median value in Fig. 5d. Upon fixation (Fig S9a), peroxisomes show less variable pH and lose the ln(G/R) dependence on G intensity seen in live cells (Fig S9b). This implies that the ln(G/R) dependence on G intensity in live cells arise from the actual pH variation of peroxisomes in live cells, and that the higher variability of ln(G/R) reports the heterogeneity in peroxisomal pH.

**Supplementary Note 14. Inter-peroxisomal pH variance validation via quantitative analysis**

To cross-verify whether the observed pH differences between peroxisomes in a single cell are the results of imaging artifacts, we fixed and permeabilized cells with methanol after transfection with HT7-SKL and SeRapHin staining as described in **Supplementary Note 11**. The obtained G/R heatmaps show decreased variance by methanol fixation as shown in **Fig. 5 b,c**. Further validation via quantitative approach was done as following. From segmented, max. projected R image, single peroxisome r.o.i.s are obtained by Fiji plugin 'Trackmate' [3]. From the r.o.i.s, ln(G/R) value of each peroxisome was calculated. ln(G/R) was selected over G/R to avoid variance scaling by the magnitude of G/R. The ln(G/R) values from peroxisomes with SeRapHin G fluorescence intensity over 2000 were selected to avoid variance skew by low signal to noise ratio, and standard deviation of ln(G/R) was compared between live and fixed cells. The standard deviation, or variance of the SeRapHin readout significantly decrease with fixation, validating that the observed variance of SeRapHin readout means inter-peroxisomal pH difference.

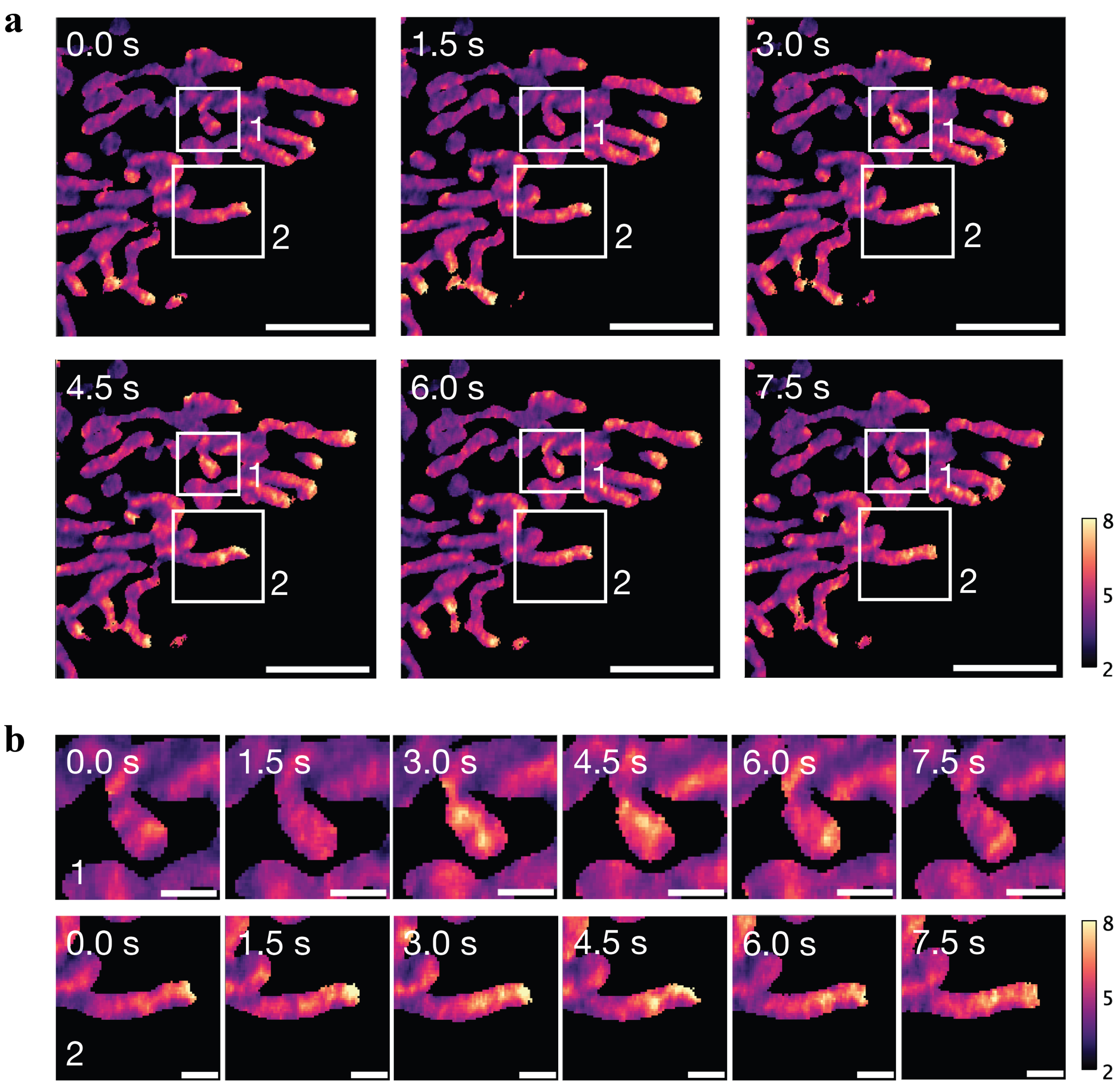

**Supplementary Figure S10. Intra-mitochondrial matrix SeRapHin R/G gradients show continuous patterns over time a**, R/G timelapse image of SeRapHin localized to mitochondrial matrix with the interval of 1.5s. Boxes 1 – 2 are shown in b. Scale bar = 5 μm **b**, Zoomed in images of boxes 1 – 2 in a. Box 1 shows a rise and decay of high R/G region in a mitochondrion, Box 2 shows a sustained intra-mitochondrial matrix gradient with higher R/G towards the right end of the mitochondrion. Scale bar = 1 μm

**Supplementary Note 15. Continuous SeRapHin R/G patterns over time further validates the Intra-mitochondrial matrix pH gradients (for Supplementary Figure S10)**

We observed the time-dependent behavior of the intra-mitochondrial gradient. Hela cells transiently expressing mitomatrix-HT7 were labeled with SeRapHin as in **Supplementary Note 10** and imaged under confocal microscope. A single-plane image was taken with 1.5s time inverval. Image analysis was done following **Extended Fig. 1**, to obtain R/G image for each timepoint. To minimize the effect of low signal to noise ratio regions, pixels with G fluorescence intensity smaller than 2000 are excluded in the final R/G map. If the gradients in R/G arise due to random noise, the high R/G spots will have random localization, and the high R/G spot's lifetime at a specific region will not exceed a single timeframe. The obtained time dependent R/G map (**Supplementary Fig. S10**) show that the high R/G spots are persistent, spanning several timepoints, reaffirming that the observed intra-mitochondrial matrix R/G differences are not due to random noise. Also, the different lifetimes of the R/G gradients point towards the temporal variability of the matrix pH gradients.

**Supplementary Note 16. Intra-mitochondrial pH gradient validation via variance quantification**

To cross-verify whether the observed pH gradient within mitochondrial matrix is the result of an imaging artifact, we collapsed any likely mitochondrial pH gradient by fixing HeLa cells expressing Mitomatrix-HT7. Since we used the same imaging conditions for live and fixed cells, if methanol fixation homogenizes the R/G signal from SeRapHin, it follows that R/G variations observed, if any, in live cells correspond to genuine variations in matrix pH. Methanol fixation was done as described in **Supplementary Note 13**. The data in **Fig. 5** shows much less variation in the R/G heatmaps of fixed cells compared to live cells. The variance was quantified as follows. The SeRapHin labeled mitochondrial images of live and fixed cells were obtained retaining the same pixel size. By the method described in **Extended Fig. 1**, the segmented and max. z-projected R and G images are obtained. Only the cells with mean G fluorescence intensity that exceeds 2000 are selected to avoid any variance arising from to low signal to noise ratio (SNR). The R and G images were mean filtered (kernel size=1, NaN pixels are excluded from mean calculation) and R/G ratio image was obtained. To avoid the effect on the variance scaling by the R/G value, ln(R/G) image was obtained for comparison between fixed and live mitochondria. After mean filtering to minimize the effect of salt & pepper noise (kernel size=3 NaN pixels excluded from calculation), the ratio of outlier pixel number from the total mitochondrial pixel number was calculated. The outlier pixels are defined as pixels with ln(R/G) values higher than the mean ln(R/G) + 0.2 of the images. Fixed cells had significantly lower ratio of high ln(R/G) outliers compared to live cells, reaffirming the observed intra-mitochondrial matrix pH gradient.

**Supplementary Note 17. Intra-ER pH gradient validation via variance quantification**

To test whether the observed pH differences within the ER are genuine and depend on the V-ATPase activity, we inhibited V-ATPase with Bafilomycin A1, constructed G/R heatmaps and compared them to that of vehicle-treated cells. Since we applied the same imaging conditions for both Baf A1 or DMSO treated cells, if the G/R spread collapses in Baf A1 treated cells, it implies that variations if any in the G/R readout of live cells reflects a genuine pH variation and is dependent on the V-ATPase activity. Cells were treated as described in **Supplementary Note 13**. ln(G/R) images wer obtained following the method described in **Supplementary Note 15**, only exchanging the position of G and R images. The outlier pixels are defined as pixels with higher ln(G/R) value than the mean ln(G/R) + 0.2 of the images, and ratio of the number of outlier pixels to the total ER pixels were calculated and compared between vehicle or Baf A1 treated cells. Baf A1 treated had significantly lower outlier ratio, reaffirming that the observed intra-ER pH gradient is genuine and depends on V-ATPase activity.
